# Evidence for chronological diversification of spinal neuron subtypes by a shared sequence of transcription factors

**DOI:** 10.1101/2025.05.27.656404

**Authors:** Laia Caudet-Segarra, Kevin Sangster, Julia Sagner, Artur Kania, Andreas Sagner

## Abstract

The mechanisms underlying the generation of the immense diversity of neuronal cell types remains a fundamental question of developmental biology. In the spinal cord, different “cardinal classes” of neurons that share a common molecular identity are produced from spatially segregated progenitor domains. Within many such classes, a stereotyped sequence of divergent neuronal types of related function is generated over time, raising the question of the molecular mechanisms that control this process. Here, we show that the successive expression of mouse transcription factors Onecut2, Pou2f2 and Pou3f1 within the cardinal classes giving rise to motor and sensory circuits, correlates with the emergence of sequentially generated subpopulations of neurons within those domains. We demonstrate that the genetic loss of Pou2f2 results in impaired development of two early-born motor neuron columns and re-specification of anterolateral system projection neurons as a later-born subset. Similarly, we show that Pou3f1 expression is required for the normal development of later-born subsets of motor neurons and anterolateral system projection neurons. Together, our observations provide functional evidence that horologic diversification of output neurons of spinal motor and sensory circuits are driven by a conserved sequential order of expression of transcription factors.

## Introduction

The vertebrate nervous system is the most complex organ system known. Its intricacy is made evident by the astonishing number of neuronal identities based on anatomical, morphological, functional and molecular criteria. The spinal cord stands out as one of the best molecularly and functionally characterized regions of the nervous system. It is a hub for the integration, gating, and relay of sensory information from the periphery to the brain and for the execution of motor commands transmitted from the brain to motor neurons (MNs) that control muscles in the body ^1–3^. Spinal neurons sharing a particular function originate from spatially-restricted populations of progenitors, but recent evidence suggests that the time of spinal neuron birth may further diversify them through a common horological program.

Spatial patterning of the spinal cord starts with opposing morphogen gradients partitioning neural progenitors into different domains arrayed along the dorsal-ventral (DV) axis. These give rise to neurons with distinct cardinal identities aligning with the broad sensory-motor division of labour ^1,4,5^. The diversity of motor neurons (MNs) derived from a ventral cardinal domain, reflects the location of their target muscles. Thus, lateral motor column (LMC) MNs innervate limb-level muscles and are subdivided into medial (LMCm) and lateral (LMCl) divisions, innervating ventral and dorsal limb muscles respectively ^6,7^. Non-limb MNs are divided among the medial motor column (MMC) innervating axial muscles, and the hypaxial motor column (HMC), innervating hypaxial muscles. Preganglionic column (PGC) MNs mediate autonomic functions by innervating sympathetic chain ganglia. The intersection of Hox-class, Foxp1, LIM homeobox, and other transcription factors (TFs) endows MNs with a columnar identity reflecting their rostrocaudal and divisional location ^8–11^. While MN birth-dating studies suggest that the development of motor columns follows pre-defined timelines, whether their diversification mechanisms have a chronological component is less clear.

Another highly diverse neuronal population of spinal projection neurons is at the origin of the anterolateral system (AS), tasked with conveying pain, temperature, and crude touch information to the brain. AS neurons largely originate from the dorsal dI5 spinal cord domain and can be divided into two broad groups: AS neuron subpopulations in the superficial layers of the dorsal horn have small receptive fields, are tuned to specific stimuli and are thought to relay the discriminatory component of their corresponding somatosensory modality, such as stimulus location or identity ^12–14^. In contrast, AS neuron subtypes in deeper laminae of the dorsal horn are tuned to a variety of stimuli and innervate brain structures that appraise the motivational dimension of sensory stimuli ^15^. A substantial proportion of superficial and deep AS neurons express the TF Phox2a which is essential for their normal molecular specification, connectivity and nociceptive function ^16^. The spatial distribution and stereotyped birth order of Phox2a-expressing embryonic AS neurons allows their division into the precursors of lamina I neurons (Phox2a^SUP^) and two subpopulations of precursors of deeper neurons (Phox2a^Deep^) ^16^. Besides the requirement of Phox2a for the molecular specification, connectivity and nociceptive function of AS neurons, there are few insights into the molecular mechanisms that control their development. Their prefigured birth order raises the possibility that chronological diversification programs could play a role in their ontogeny.

Classical studies in *Drosophila* and more recent experiments in vertebrates provide insights into the developmental logic of temporal specification of neurons by a stereotyped procession of TFs in their progenitors or stem cells ^17,18^. Such temporal TFs instruct the stem cell to generate a particular neuronal type, until its expression is exchanged with the next TF in the sequence, instructing the generation of a different neuronal type ^17,19–23^. Examples of temporal TFs consistent with this model in the context of retinal neuron development include Ikzf1, Ikzf4, Foxn4, Casz1, Onecut2 (OC2), Pou2f2, and Nfib ^24–29^. In the spinal cord, timing of birth appears to be a crucial factor influencing the diversification of a variety of neurons ^30–33^, including MNs among which early-born LMCm neurons regulate the specification of later-born LMCl neurons ^34,35^. Recent single cell RNA sequencing (scRNAseq) analyses suggest that temporal TFs such as OC2, Pou2f2, Zfhx3, Nfib and Neurod2 are expressed in a conserved chronological sequence correlating with the birth time of most cardinal classes in the mouse and human spinal cord ^36–39^. However, there remains a significant gap in linking hallmarks such as combinatorial expression of distinct spatial and temporal TFs to previously defined molecular, anatomical, or physiological neuronal subtypes. More importantly, functional studies examining the role of these TFs in the spinal cord, other than those demonstrating the requirement of Onecut1 and OC2 for the generation of MN diversity and the generation of Renshaw neurons ^32,40^, have yet to be conducted.

Here we provide evidence that a temporal TF sequence —OC2 > Pou2f2 > Pou3f1 (OPP) — organises spinal neurons within multiple cardinal domains according to their birth chronology. The expression of OPP TFs also correlates with distinct functional subtype identities within two cardinal domains with long-projecting neurons: dividing MNs into distinct motor columns, and AS neurons into superficial versus deep laminar subtypes. Evidence from genetic perturbations of Pou2f2 and Pou3f1 expression argues that these are required for the normal molecular specification of motor and AS neuron subtypes. Together, our study links the expression of temporal TFs in different cardinal domains to the specification of neuronal subtype identities, and provides evidence that the diversification of neuronal cell types emerges through the integration of shared spatial and temporal patterning programs in the embryonic spinal cord.

## Results

### A chronological order of expression of OC2, Pou2f2 and Pou3f1 TFs in the embryonic spinal cord

Past spinal cord birthdating experiments have suggested two general chronological principles: (1) a ventral to dorsal cardinal group birth progression ^6,41^, and (2) the birth of projection neurons preceding the birth of local interneurons within a cardinal group ^37^. Recent transcriptomic analyses reveal a small set of mRNAs encoding TFs OC2, Pou2f2, Zfhx3/4, Nfib and Neurod2/6 correlates with neuronal birth times within most cardinal domains of spinal neurons (Fig. 1A) ^36,37^. We revisited these studies by re-analysing single spinal cord cell transcriptomic data to reveal neuronal TF expression differences between embryonic day (e) 9.5 and e13.5 (Fig. S1A-D) ^36^. Besides providing further support for the temporal sequence of OC2 followed by (>) Pou2f2 and Zfhx3/4 > Nfib and Neurod2/6, our re-assessment suggested that the timing of expression of Pou3f1 (also known as Scip or Oct6) may fit this chronology (Figs. 1B and S1C, D).

**Figure 1:**
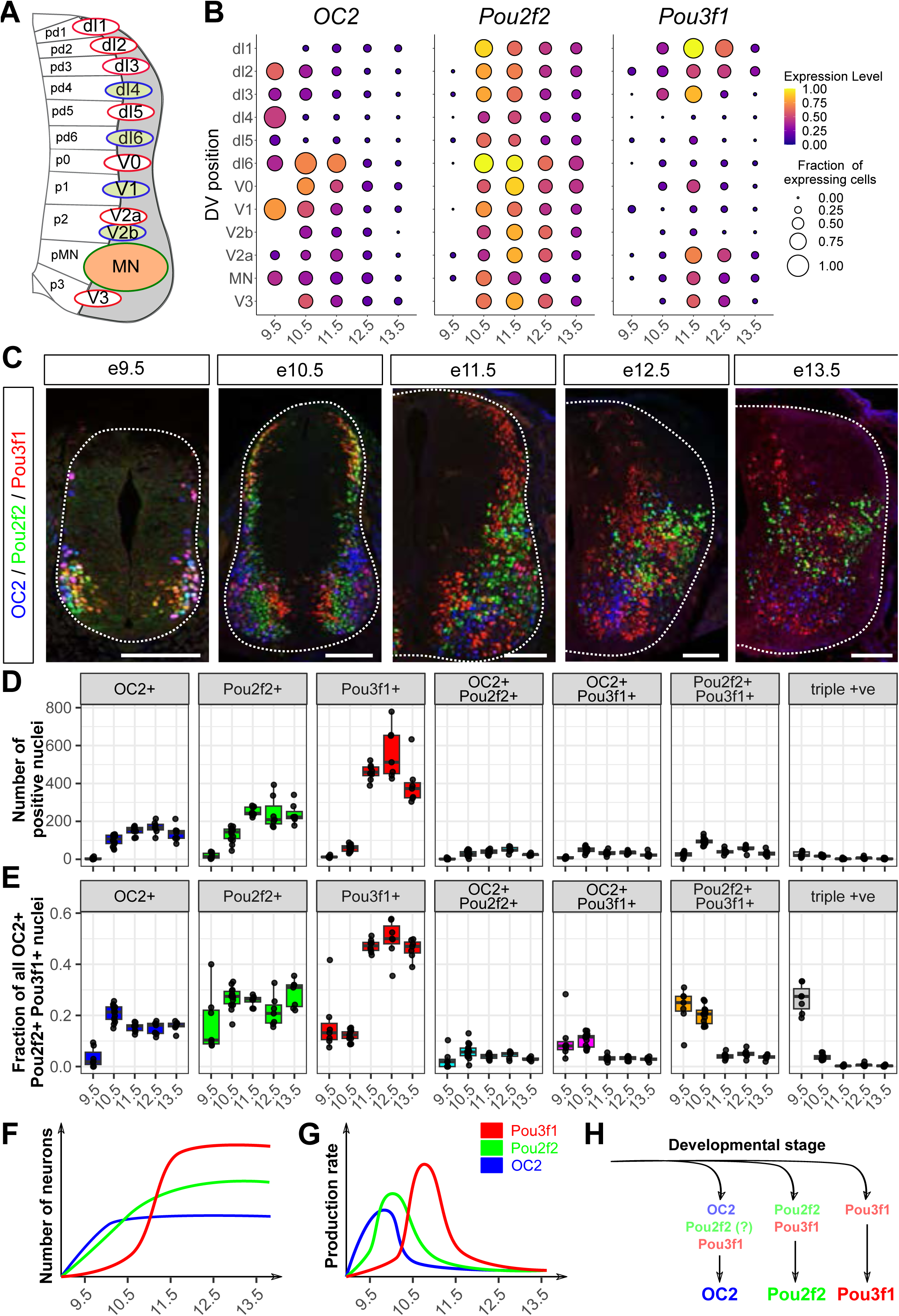
Characterization of OPP expression dynamics. (A) Dorsal-ventral patterning partitions the spinal cord into distinct progenitor domains that subsequently generate distinct classes of neurons (B) Spatio-temporal RNA expression dynamics of OPP mRNAs in scRNAseq data from Delile et al. 2019. (C) OPP protein expression dynamics in e9.5 – e13.5 cervical spinal cord cryosections. (D, E) Quantification of the absolute number (D) and relative fraction of all OPP+ nuclei (E) that express OPP TF combinations (F, G) Scheme indicating the increase in absolute numbers (F) and inferred production rates (G) of OPP neurons (H) Model for OPP neuron generation. OPP single-positive neurons are sequentially generated during development. However, early-born neurons pass through a state where they co-express other OPP TFs before consolidation to a single OPP TF identity Scale bars in C = 100 µm

Fragmentary evidence suggests that protein expression of OC2, Pou2f2 and Pou3f1 (OPP) TFs is also temporally ordered within different cardinal groups ^42–44^. To test this possibility, we examined their protein expression between embryonic day (e) 9.5, the onset of birth of projection neurons and ventral neuronal types, but with a negligible population of interneurons, and e13.5, the time of birth of many dorsal interneurons. At e9.5 OPP expression was concentrated in the ventral spinal cord, the location of birth of motor neurons, and within a limited number of dorsal neurons (Fig. 1C). At subsequent ages, we found a dorsal expansion of the OPP-expressing (+) cells, co-incident with the shift of the principal site of neurogenesis from the ventral to the dorsal spinal cord. Over the ages examined, the number of cells that were OC2-only (not expressing Pou3f1 or Pou2f2) increased between e9.5 and e10.5, the number of Pou2f2-only increased between e9.5 and e11.5, while the number of Pou3f1-only cells was initially low, but then increased sharply between e10.5 and e11.5 (Fig. 1D, E). The number of cells expressing each TF stayed roughly constant after e11.5, suggesting that most cells maintained expression of OPP TFs over the ages analysed. Together, these data suggest that the expression of OPP proteins follows a chronology suggested by their mRNA expression, with OC2 and Pou2f2 expression preceding that of Pou3f1 (Fig. 1F-H).

Our analysis revealed a significant fraction of spinal cord cells that co-expressed two or three OPP proteins at early ages. In contrast, at later times most OPP+ cells expressed either OC2, Pou2f2 or Pou3f1 (Figs. 1D-E and S1E-G), raising the possibility that cells expressing multiple OPP TFs, eventually express only one. In support of this, most cells in the dorsal third of the spinal cord, presumably dI1, dI2 and dI3 neurons that eventually migrate ventrally, co-expressed Pou2f2 and Pou3f1 at e10.5 (Fig. 1C). In contrast, at e11.5, the number and fraction of Pou2f2+ Pou3f1+ cells decreased and most cells in the dorsal third of the spinal cord were now only expressing Pou3f1, while Pou2f2-only cells were largely located in the ventral and intermediate locations (Fig. 1C). To gain further insight into OPP expression dynamics, we compared OPP TF expression in neuronal progenitors and maturing post-mitotic neurons between e9.5 and e11.5 by using the neurogenic marker Neurogenin2 (Neurog2) (Fig. S2A, B) ^45,46^. At e9.5, approximately half of Pou2f2+ and Pou3f1+ (single TF expressing) and Pou2f2+ Pou3f1+ cells expressed Neurog2 (Fig. S2A-C). At subsequent ages, with the decrease in Neurog2 expression as neural precursor exit from mitosis shifts from ventral to dorsal domains, the proportion of OPP+ cells that were also Neurog2+ waned (Fig. S2A-C). Between e9.5 and e10.5, expression of Neurog2 in Pou2f2 single-positive cells was extinguished, while the fraction of Pou2f2+ Pou3f1+ cells decreased and the fraction of Pou3f1-only increased. Finally, at e11.5, little co-expression between Neurog2 and OPP TFs was detected throughout the spinal cord (Fig. S2A-C). Thus, while at early ages a significant fraction of neurogenic progenitors expresses OPP proteins, at later stages, their expression shifts to postmitotic neurons.

We next aimed to test if all or only a subset of neurons expressing a specific OPP TF transiently co-expresses the other OPP TFs. To this end, we performed short-term lineage-tracing of OC2+ neurons using a *lacZ* transgene expressed from the endogenous *Onecut2* locus ^47^, and quantified the fraction of lacZ+ OPP-expressing neurons at e11.5 (Fig. S2D-F). Approximately 85% of the OC2+ neurons, and only ∼18% of Pou2f2 and ∼3% of Pou3f1-positive neurons were lacZ+ (Fig. S2F). Similarly, most lacZ+ neurons expressed OC2, either alone (∼42% of lacZ+ neurons) or in combination with Pou2f2 (∼20%). In contrast, only a small fraction of lacZ+ neurons expressed Pou2f2-only (∼18%) or Pou3f1-only (∼4%) (Fig. S2E). Thus, most neurons that initiate OC2 expression, as either OC2+ only or in combination with Pou2f2 and/or Pou3f1, acquire an OC2+ identity by e11.5, while most Pou2f2 and Pou3f1 single-positive neurons detected at this stage did not pass through a OC2+ state. Taken together, these observations support a model in which a large fraction of early-generated spinal neurons transiently co-expresses stage-specific combinations of OPP TFs during or shortly after neurogenesis, followed by exclusive expression of one of the OPP TFs (Fig 1H).

### Complementary expression of OC2, Pou2f2 and Pou3f1 within individual spinal cardinal domains

Next, we examined whether the above relative abundance of OC2, Pou2f2 and Pou3f1-expressing neurons was preserved within the different cardinal domains at e11.5. We thus examined the expression of OPP proteins in combination with the dI3 marker Isl1/2 (dorsally located), the dI5 marker Lmx1b, the V0 marker Evx1, the V2a marker Vsx2, the MN marker Isl1/2 (ventrally located) and the V3 marker Nkx2.2 (Figs. 2A and S3A). These experiments revealed that indeed, each of these cardinal domains contained OC2, Pou2f2 and Pou3f1-expressing neurons, and many of these expressed only one of the OPP TFs. The number and fraction of neurons expressing a single OPP protein differed between the cardinal domains examined (Figs. 2B, C). The number of early-born OC2 and Pou2f2-single positive neurons was for example higher in MNs, while the fraction of Pou3f1+ neurons was higher in dI5 and dI3 neurons compared to MNs. These observations are consistent with the increased neurogenesis rate in the ventral spinal cord, especially of MNs, and the comparatively lower neuronal differentiation rate in the dorsal part of the spinal cord at early ages ^48–50^. Moreover, while some cardinal domains like V3 or V0 could be parcellated into neurons expressing a single OPP protein, others, such as V2a and motor neurons contained a greater fraction of neurons expressing at least two different OPP proteins. (Figs. 2C and S3B-D). Together, these observations suggest that OPP expression segregates spinal cord cardinal domains into molecularly distinct subtypes of neurons, and that the relative abundance of these subtypes varies between cardinal classes, largely in accordance with previously observed macroscopic spatial patterns of progenitor exit and neuronal differentiation.

**Figure 2:**
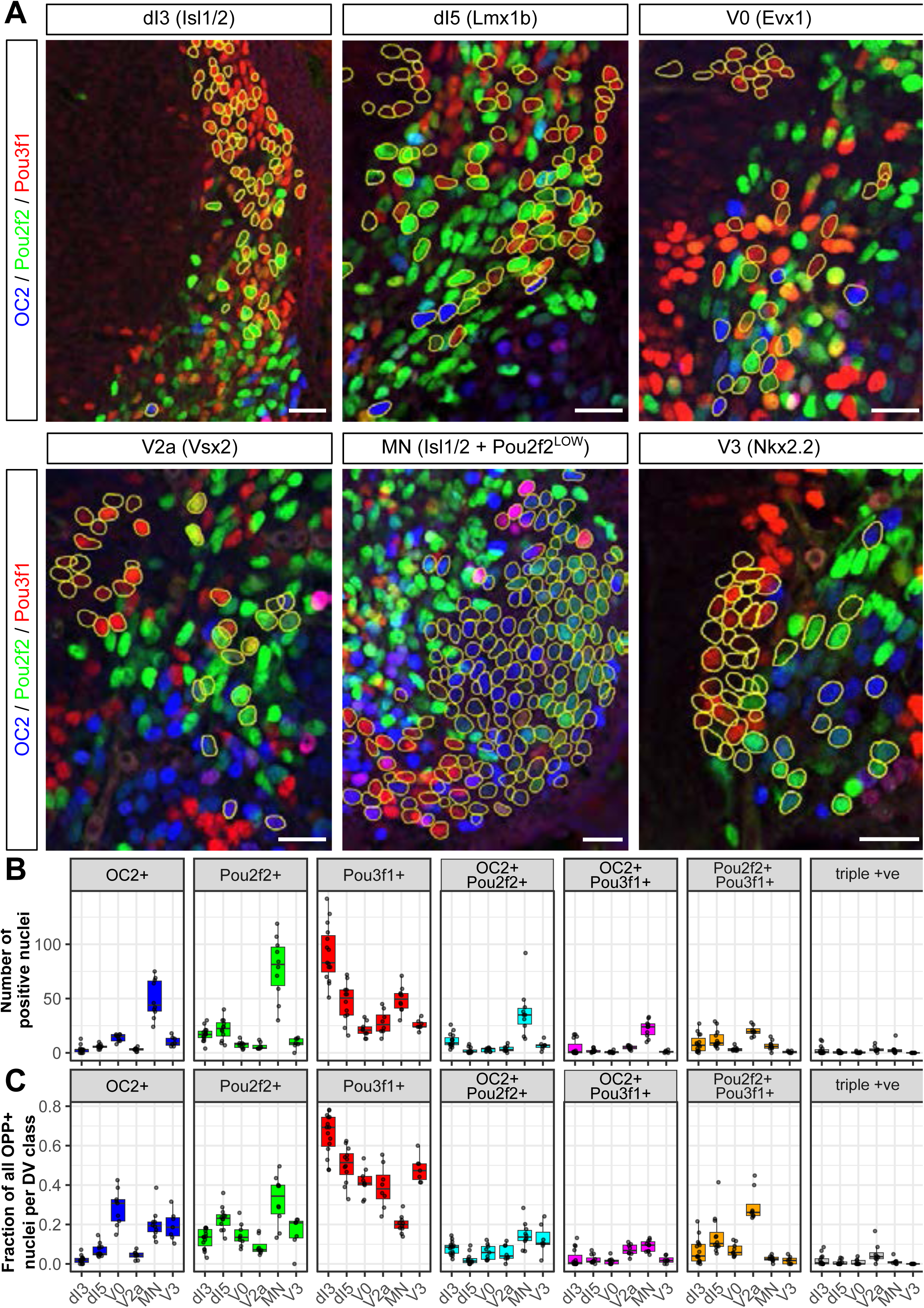
OPP TFs parcellate spinal neuronal classes into distinct subtype identities. (A) OPP protein expression in different DV classes of neurons in e11.5 cervical spinal cord sections. Segmentation of neurons with specific DV identities (yellow outlines) were obtained from co-staining with the indicated DV markers (see Fig. S3A). (B, C) Quantification of the absolute number (B) and relative fraction of all OPP+ neurons (C) in the indicated DV classes of neurons Scale bars in A = 100 µm

Previous analyses identified Pou3f1 as differentially expressed between neurons belonging to the different cardinal classes ^36^ and comparison of OPP spatiotemporal expression dynamics suggested low Pou3f1 levels in inhibitory dI4, dI6, V1 and V2b neurons (Figs. 1B and S4A,B). To directly test this prediction, we stained e11.5 cryosections for Pou3f1 in combination with Pax2, which labels inhibitory dI4, dI6 and V1 neurons ^51^, and the intermediate dorsal marker Lbx1, which labels dI4-6 neurons ^52,53^ (Fig. S4A-C). Consistent with the predictions of the scRNAseq data, no Pou3f1 expression was detected in Pax2+ Lbx1+ dorsal and Pax2+ Lbx1-ventral inhibitory neurons (Fig. S4C, D). Thus, unlike OC2 and Pou2f2, which are expressed in most cardinal classes and for which expression in dorsal and ventral inhibitory neurons has been previously documented ^32,44,54^, Pou3f1 expression is limited to cholinergic MNs and glutamatergic dI1-3, dI5, V0, V2a and V3 neurons, suggesting that not all cardinal domains can be parcellated by OPP TF expression.

### Sequential expression of OC2, Pou2f2 and Pou3f1 correlates with spinal neuron birth order

Most cardinal neuronal subtypes undergo stereotypical migrations such that their relative order of birth correlates with the location along their pre-determined route ^3^. For example, neurons from most dorsal domains migrate ventrally along commissural axons, V3 neurons migrate dorsally and some MNs follow an inside out pattern within the ventral horn ^30,34,55^. We reasoned that a temporal sequence of OPP expression within a particular group of neurons may be reflected in their spatial distribution, with earlier-born neurons being further along their usual migration trajectory compared to later born neurons. We thus determined the relative position of OC2+, Pou2f2+ and Pou3f1+ neurons within the different cardinal classes in the brachial spinal cord at e11.5 and found that it was significantly different for the populations assayed (dI3, dI5, V0, V2a, MN and V3; Figs. 3A and S4E). In agreement with the known medial to lateral or medial to ventro-lateral trajectories of dI3, V0, V2a and V3, OC2+ neurons were typically located farthest away from their birth location, Pou3f1+ neurons were closest, and Pou2f2 occupied an intermediate position. Within the MN domain, Pou2f2+ neurons were farthest, Pou3f1 closest and OC2 occupied an intermediate position, also consistent with the known inside out migration pattern of LMC MNs (see below). Together, these data are consistent with the hypothesis that OPP protein expression correlates with spinal neuron birth order.

**Figure 3:**
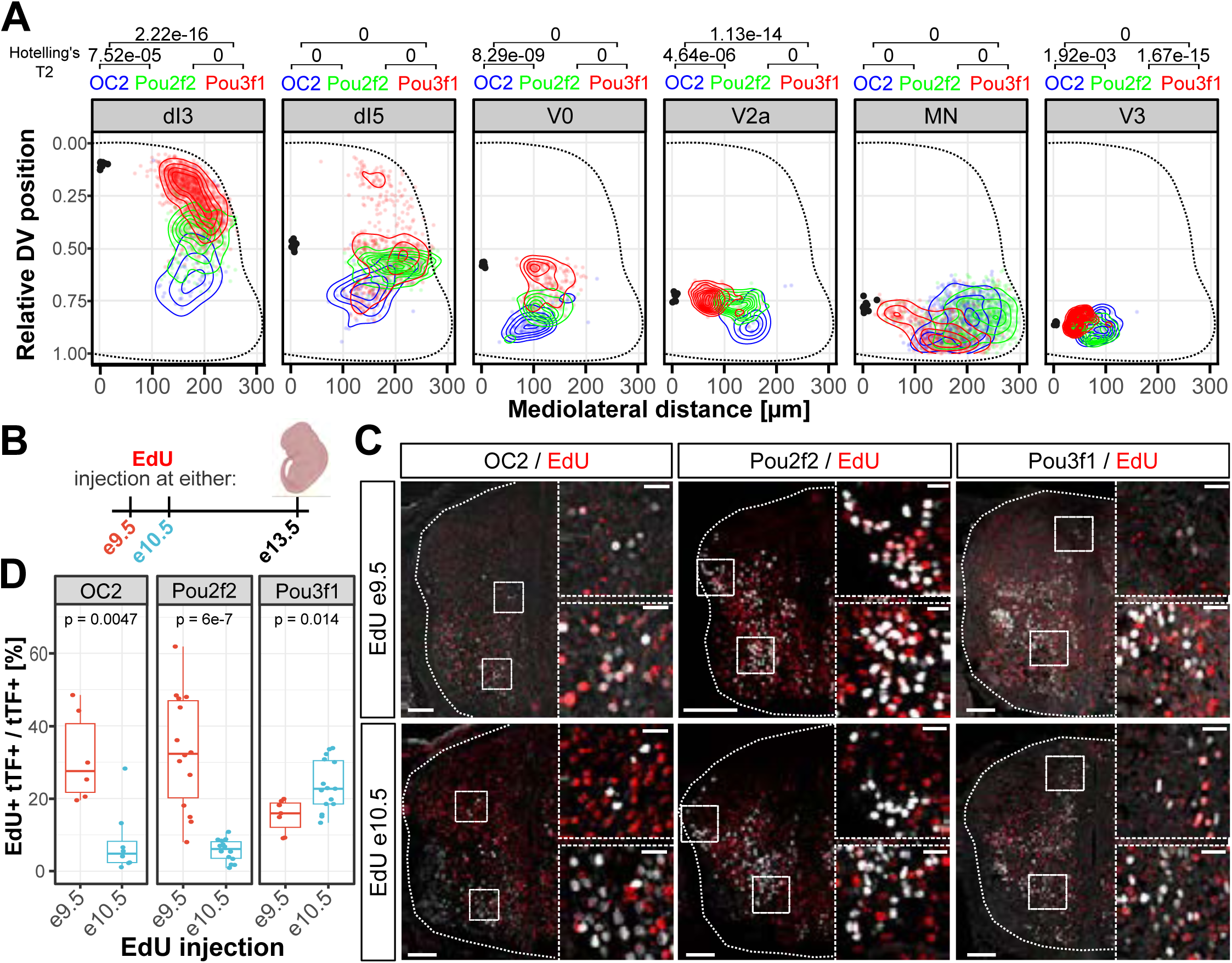
Sequential generation of OPP neurons. (A) Spatial distribution of OPP neuron subsets in most DV classes of neurons is consistent with the sequential generation of these neurons (B) Scheme outlining EdU-birth-dating experiments. EdU was injected either at e9.5 or e10.5 and incorporation into OPP subsets was analysed at e13.5. (C) OPP protein staining in combination with EdU (red). Onecut2 and Pou2f2+ neurons incorporated EdU only, when administered at e9.5 but not at e10.5. Pou3f1+ neurons incorporated EdU administered at both timepoints (D) Quantification of the fraction of OPP neurons positive for EdU. Scale bars in D = 100 µm, 25 µm

To test this hypothesis, we injected pregnant female mice at e9.5 or e10.5 with EdU, and analysed the co-incidence of OC2, Pou2f2 and Pou3f1 expression and EdU at e13.5, across all spinal cord cardinal domains (Fig. 3B). This analysis revealed that while nearly 40% of OC2+ or Pou2f2+ neurons were labelled with the EdU injection at e9.5, essentially none of them were labelled by the e10.5 EdU injection. In contrast, although ∼15% of Pou3f1+ neurons were labelled by EdU injection at e9.5, ∼25% of them were labelled by the EdU injection at e10.5 (Fig. 3C, D), indicating that while some Pou3f1+ neurons are born at the same time as OC2+ and Pou2f2+ neurons, a subpopulation of them is born at a later time. To generate a more detailed snapshot of spinal cord birth order, we injected EdU at e9.5, and then BrdU at e10.5 in the same pregnant females (Fig. S5A, B). Analysis at e11.5 revealed that Pou2f2-expressing neurons were stronger labelled by EdU (30%) than by BrdU (13%) (Fig. S5C-E). In contrast, only 17% of e11.5 Pou3f1-expressing neurons were labelled by EdU and 35% of them were labelled with BrdU, suggesting that most Pou2f2+ neurons are born before Pou3f1+ neurons (Fig. S4C-E). Together, these data suggest that the OC2 precedes (>) Pou2f2 > Pou3f1 chronological sequence of expression may be a general feature of developing spinal neurons.

### OC2 > Pou2f2 > Pou3f1 expression and motor column birth order

To determine with more precision whether a temporal TF expression sequence correlates with the sequential birth of spinal neurons, we turned to MNs, a cardinal spinal cord domain extensively classified into molecularly-distinct neuronal subtypes ^4,56^ (Fig. 4A). Past studies, whose resolution is limited by slow birthdating tracer diffusion, suggest the following order of MN birth at limb levels: LMCm birth precedes LMCl birth, which precedes MMC birth (LMCm > LMCl > MMC) ^34,35,57^. The birth order at thoracic levels is less-well resolved, although previous observations in stem cell-based differentiations suggested that HMC precedes MMC birth ^58^. We first examined the expression of *OC2*, *Pou2f2* and *Pou3f1* mRNAs in MNs identified in mouse spinal cord scRNAseq datasets from e10.5 to e12.5 ^36,59^. MN subpopulations can be molecularly distinguished based on the combinatorial expression of the TFs *Isl1*, *Isl2*, *Lhx1* and *Foxp1* and the retinoic acid synthesizing enzyme *Aldh1a2* ^8,10,11^. Moreover, all somatic MNs transiently express *Isl1* and *Lhx3* after neurogenesis, while *Lhx3* expression is maintained in MMC neurons at later stages ^11,60^. UMAP-based dimensionality reduction at e10.5 revealed that MNs from cervical and thoracic levels can be arranged along a continuous trajectory from *Olig2*/*Neurog2*-expressing pMN progenitors to MNs expressing maturing markers (Fig. S6A-C). *OC2+* cells were found in the most mature region of the differentiation trajectory containing LMCm neurons (*Isl1+, Foxp1+, Aldh1a2+*) and a cluster of neurons expressing the MMC marker *Mecom* (Figs. 4B and S6A-C). *Pou2f2* mRNA was enriched in cells at intermediate stages of the trajectory that expressed LMCl hallmarks (low *Isl1* levels, *Isl2+*, *Lhx1*+), while *Pou3f1* mRNA was enriched in *Lhx3*/*Lhx4*+ *Mnx1+* newborn MNs (Figs. 4B and S6A-C). For the e11.5 and e12.5 datasets, motor column identity could be assigned to cells based on their transcriptome profiles (Fig. S6D, E). At both ages, *OC2* mRNA was enriched in the LMCm, *Pou2f2* mRNA in the LMCl and PGC and *Pou3f1* mRNA in the MMC (Fig. S6D-F).

**Figure 4:**
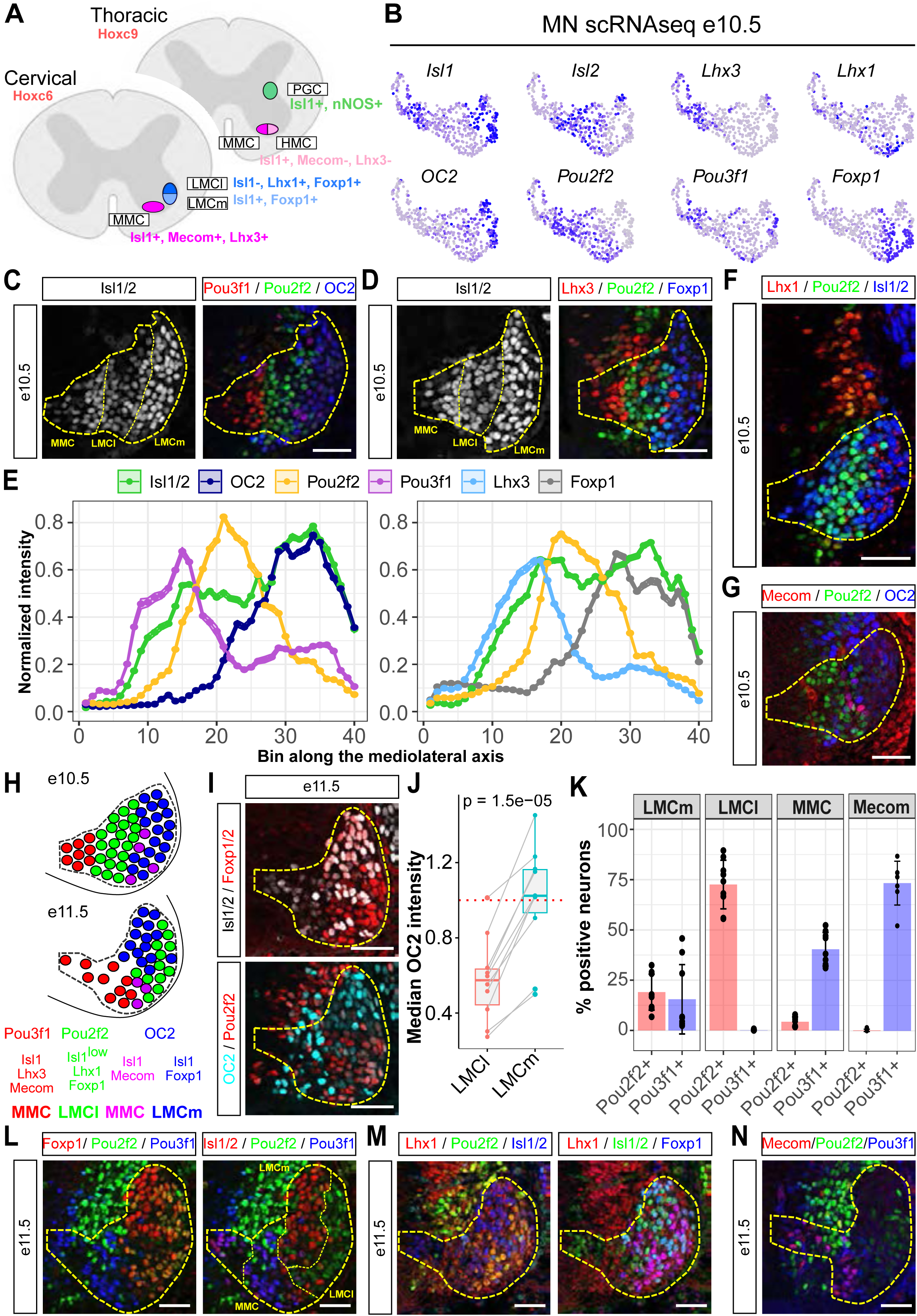
OPP expression correlates with motor column identity. (A) Cervical and thoracic MNs are partitioned into distinct motor columns that express different marker genes (B) OPP mRNA expression relative to motor column markers *Isl1*, *Isl2*, *Lhx3*, *Lhx1* and *Foxp1* in MN scRNAseq data from Delile et al. 2019. Expression of additional marker genes is shown in Fig. S6C. (C) OPP expression in MNs at e10.5. Consistent with their sequential generation, OPP+ MNs are distributed in stripes along the mediolateral axis of the spinal cord at e10.5. (D) Staining for the MMC marker Lhx3 (red), and the LMC marker Foxp1 (blue) and the temporal TF Pou2f2 (green) in MNs identified by Isl1/2 staining at e10.5. (E) Intensity distribution of OPP TFs and motor column markers (Isl1/2, Lhx3, Foxp1) along the mediolateral axis in motor neurons at e10.5. OC2 correlates with Foxp1 expression and lateral upregulation of Isl1/2 indicative of an LMCm identity. Pou2f2 expression correlates with a stripe of reduced Isl1/2 staining, indicative of the LMCl, while Pou3f1 expression correlates with Lhx3 expression in the most medial population of MNs. (F) Pou2f2 expression precedes expression of the LMCl marker Lhx1 in MNs. (G) Expression of the MMC marker Mecom is limited to few Onecut2+ MNs at e10.5. (H) Summary of OPP expression in cervical MNs at e10.5 and e11.5. OC2 neurons largely contribute to the LMCm and a small subset of MMC MNs, Pou2f2 to the LMCl, and Pou3f1-positive neurons to the MMC. (I) OC2 and Pou2f2 expression partitions the LMC into LMCm and LMCl. OC2 expression in Foxp1+ LMC neurons correlates with expression of the LMCm marker Isl1/2. Pou2f2 expression correlates with down-regulation of Isl1/2 and LMCl identity. (J) OC2 levels are elevated in the LMCm compared to the LMCl. (K) Quantification of the percentage of Pou2f2+ and Pou3f1+ neurons in the LMCm, LMCl, MMC (Isl1/2+ Foxp1-) and Mecom+ MMC neurons. (L - N) Pou2f2 and Pou3f1 expression in distinct motor columns. e11.5 cervical MNs were stained for Pou2f2 and Pou3f1 in combination with motor column markers. (L) Pou2f2 is expressed in Isl1/2- Foxp1+ LMCl neurons. Pou3f1 expression is limited to Isl1/2+ Foxp1-MMC neurons. (M) Co-expression between Pou2f2 and Lhx1 in Isl1/2- Foxp1+ LMCl neurons (N) Co-expression between Pou3f1 and Mecom in MMC neurons. Scale bars = 50 µm

Next, we examined the expression of OPP proteins in the cervical spinal cord at e10.5 and e11.5, when MNs undergo a complex migration. During it, late born MNs such as those of the LMCl, displace earlier born LMCm MNs to occupy the lateral-most position within the ventral horn ^11,34^. At e10.5, when MNs express Isl1 and Isl2 TFs (Isl1/2), the expression of OC2, Pou2f2 and Pou3f1 in MNs was in largely complementary stripes along the mediolateral axis (Figs. 4C-E and S7A). LMCl formed a band of low Isl1/2 expression (Isl1/2^LOW^), and expressed Pou2f2. The expression of the LMCl TF Lhx1 remained low in LMCl MNs at this time (Figs. 4F, S7C and S8C). Lateral to these were LMCm MNs with intense Isl1/2 expression (Isl1/2^high^) co-expressing the pan-LMC marker Foxp1 and OC2 (Figs. 4C-E, H and S7A, B). Pou3f1 was predominantly found in a population of MNs medial to LMC neurons, co-expressing the MMC and maturing neuron marker Lhx3. (Fig. 4C-E). We also noted a small population of OC2+ MNs lateral to the Pou2f2 stripe that also expressed Lhx3 and the MMC marker Mecom (Figs. 4D, G and S7B, D), echoing findings from the chicken spinal cord and suggesting that OC2+ neurons contribute to LMCm and MMC in mice ^61^. Despite this exception, our observations corroborate the e10.5 scRNAseq observations and correlate OC2, Pou2f2 and Pou3f1 expression with, respectively, LMCm, LMCl and MMC MN identities.

At e11.5, LMCl MNs reached their destination in the lateral LMC and continued expressing Pou2f2, while LMCm neurons were displaced medially and maintained their expression of OC2 (Fig. 4H-J, M; Fig. S7C). Pou3f1 was expressed in a large fraction of Mecom+ MMC neurons (Figs. 4K, L, N and S8B, D; ^10^). Many of them were present close to the ventricular zone containing neuronal progenitors, contrasting the lateral location of LMC neurons. This is consistent with a large fraction of MMC neurons being born at an age when the generation of LMC neurons had already stopped (Fig. 4H; Fig. S8D). In line with this hypothesis, quantification of the abundance of LMC and MMC neurons between e9.5 and e12.5 revealed that most LMC neurons are generated prior to e11.5, while MMC neuron numbers continued to increase until e12.5 (Fig. S8E, F). Remarkably, in the e11.5 thoracic spinal cord, OPP proteins were expressed in mediolateral stripes like those seen in the e10.5 cervical spinal cord. Pou2f2 was found in cells expressing the PGC marker Foxp1, while OC2 expression was lateral to these, in the presumptive HMC domain, and the MMC marker Lhx3 overlapped with Pou3f1 (Fig. S9A, B). We next examined the expression of these proteins in the thoracic spinal cord at e12.5 and e13.5, a time at which the identities of these MNs columns can be distinguished molecularly and anatomically (Fig. 4A). There, OC2 expression was largely absent, while Pou2f2 was expressed in most PGC (nNOS+) neurons but not in HMC or MMC neurons, and Pou3f1 expression was restricted to MMC neurons (Figs. S6E-G and S9C-E). Together, these observations suggest that the TFs OC2, Pou2f2 and Pou3f1 are expressed in different MN columns, and their chronological expression (OC2 > Pou2f2 > Pou3f1) aligns with the sequential generation of MN columns at limb and thoracic levels (LMCm/HMC > LMCl/PGC > MMC) (Fig. S9F).

### OC2, Pou2f2 and Pou3f1 delineate functionally distinct subpopulations of motor-related V3 neurons

We next asked whether the chronology of OC2, Pou2f2 and Pou3f1 expression can be correlated with functionally defined neuronal populations within a non-MN cardinal domain related to motor function. Neurons from the V3 cardinal group are segregated into dorsal and ventral subtypes (V3_D_ and V3_V_, respectively) which can be further subdivided into subpopulations with distinct cell morphologies, projection patterns, electrophysiological properties and recruitment during locomotor behaviours ^30,62^. To determine whether the generation of V3_V_ neurons expressing the TF Olig3 fits into the OC2 > Pou2f2 > Pou3f1 temporal progression, we examined a scRNAseq dataset from e12.5 embryos enriched for motor and V3 neurons ^59^ (Fig. S10A-D). UMAP-based dimensionality reduction and pseudotime reconstruction of the V3 neurons revealed a continuous trajectory from *Sox2* and *Neurog3* mRNA expressing neurogenic cells to those that expressed *Lhx1* mRNA, a post-mitotic V3_D_ neuron identity marker ^5,42^ (Fig. S10A-D). While such neurons expressed *OC2* and *Pou2f2* mRNA, cells in the middle of the pseudotime trajectory, thus presumably later born, expressed *Pou3f1* as well as the V3_V_ marker *Olig3* mRNA (Fig. S10A, D). In line with this, we detected a complementary expression of Pou2f2 and Pou3f1 proteins in, respectively, V3_D_ and V3_V_ domains of the e11.5 mouse spinal cord (Fig. S10E), consistent with recent work demonstrating that the V3_D_ subpopulation can be divided into early and later born neurons by their respective expression of OC2 and Pou2f2 ^63^. More generally, OPP TF expression correlates with the chronology of diversification of the V3 cardinal domain

### OC2 > Pou2f2 > Pou3f1 expression and AS projection neuron birth order

To determine whether OPP TFs could contribute to the generation of neuronal diversity in the dorsal spinal cord, we examined their expression in AS neurons defined by the embryonic expression of the TF Phox2a in dI5 cardinal domain neurons, which can be divided into three sequentially-generated populations: early Phox2a^Deep^ > Phox2a^Sup^ > late Phox2a^Deep^ ^16^ (Fig. 5A). These subpopulations can already be identified in the embryonic spinal cord based on their characteristic positions ^16,64^ (Fig. 5B). At e12.5, late Phox2a^Deep^, early Phox2a^Deep^ and Phox2a^Sup^ occupy distinct positions along the mediolateral axis at intermediate DV levels of the spinal cord. At e13.5, Phox2a^Sup^ are found ventral to the lateral-most edge of the dorsal horn, while late Phox2a^Deep^ are located dorsally near the ventricular zone (Fig. 5A, B; Osseward et al., 2021). By e15.5, Phox2a^Sup^ cells migrate tangentially into lamina I, while the late born Phox2a^Deep^ neurons migrate laterally and intermingle with early Phox2a^Deep^ neurons (Fig. 5B; Roome et al., 2020). Recent RNAseq data suggest that these three populations produce five molecularly distinct adult AS neuron populations ^65^. We reanalysed these data by asking whether ALS neuron subtypes can be differentiated via their expression of OPP mRNAs. Intriguingly, *Pou2f2* mRNAs was enriched in ALS1, ALS2 and ALS3 and *OC2* mRNA in ALS5 neurons; however, *Pou3f1* mRNA was not detectable in this dataset (Fig. 5C). This suggests that early and late Phox2a^Deep^ neurons produce ALS5 and ALS4 adult AS neurons, respectively, while Phox2a^Sup^ produce ALS1, ALS2 and ALS3 adult AS neurons.

**Figure 5:**
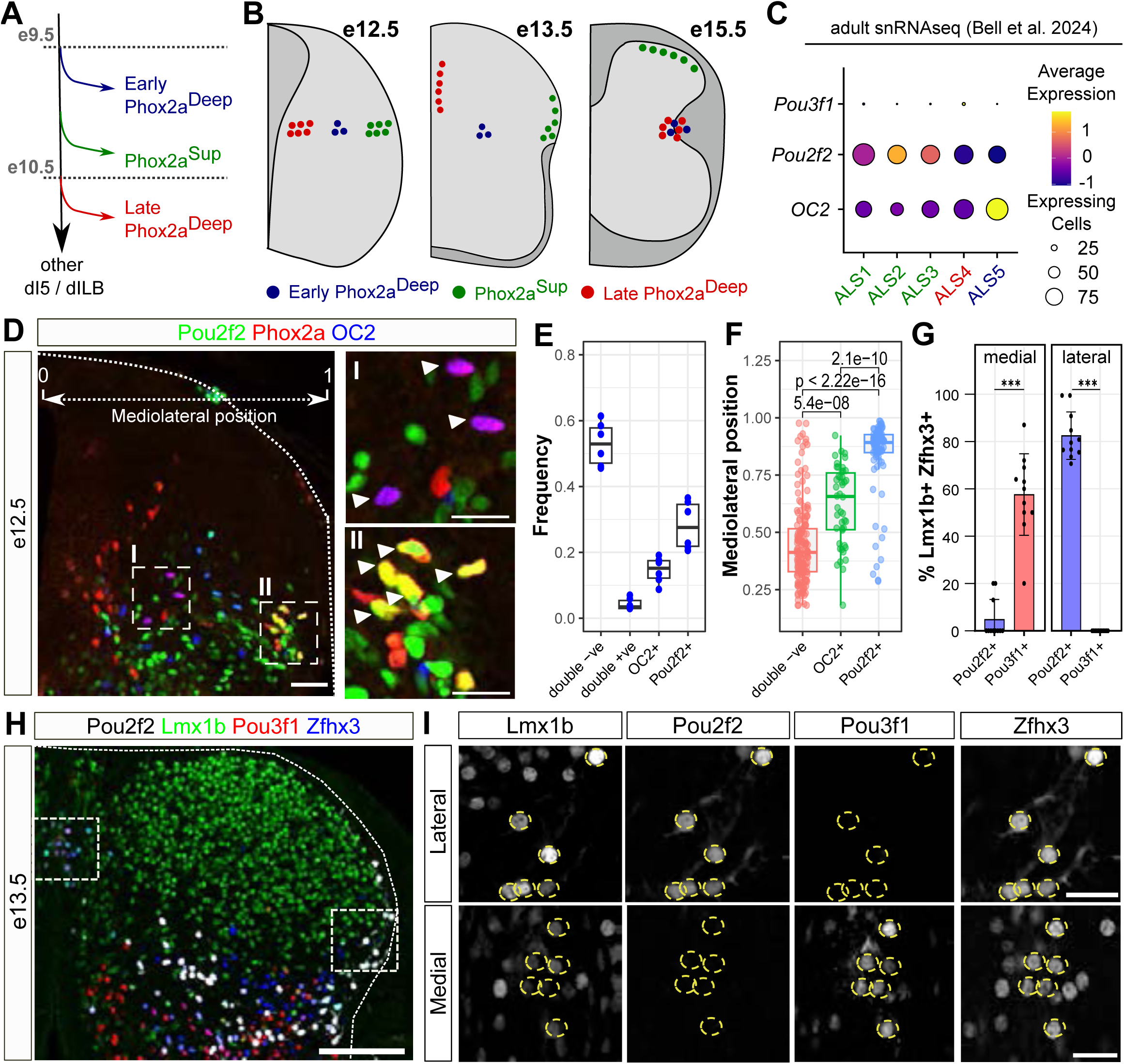
OPP expression in distinct AS neuron subtypes. (A) Sequence of AS neuron subtype generation (B) Migratory routes of AS neuron subtypes as outlined in Roome et al. 2020 and Roome et al. 2022. (C) Enrichment of *Pou2f2* in ALS1-3 lamina I Phox2a^Sup^ cells and of *OC2* in ALS5 Phox2a^Deep^ cells in adult AS neuron snRNAseq data from Bell et al. 2024. (D - F) Characterization of OC2 and Pou2f2 expression in Phox2a+ AS neurons at e12.5. (D) Consistent with their expression in Early Phox2aDeep and Phox2aSup neurons, OC2 and Pou2f2 are expressed in AS neurons localized at intermediate mediolateral (I) and lateral positions (II) respectively. (E) Fraction of OC2 Pou2f2 double-negative, double-positive and single-positive neurons Phox2a+ neurons (n = 4). (F) Quantification of the mediolateral position of OC2 Pou2f2 double-negative, OC2+ and Pou2f2+ single-positive neurons (n = 3). (G - I) Differential expression of Pou2f2 and Pou3f1 in lateral and medial Lmx1b Zfhx3 double-positive neurons at e13.5.(G) Percentage of medial and lateral Lmx1b Zfhx3 double-positive neurons expressing Pou2f2 and Pou3f1. (H) e13.5 cervical spinal cord section stained for Lmx1b, Zfhx3, Pou3f1 and Pou2f2. (I) Single channel images of the boxed regions in H. Pou3f1 staining in Lmx1b Zfhx3+ neurons is only detected in medial Late Phox2a^Deep^ and Pou2f2 staining in lateral Phox2a^Sup^ neurons. Scale bars in D, H = 50 µm; in DI, DII = 25 µm; in I = 10 µm

Could selective expression of OPP in embryonic Phox2a neurons presage the selective OPP expression in adult AS neurons? We examined Phox2a, OC2 and Pou2f2 expression in e12.5 spinal cord (Fig. 5D), and quantified the position of OC2+ Phox2a+ and Pou2f2+ Phox2a+ cells along the mediolateral axis of the spinal cord. This analysis revealed 3 populations: OC2, Pou2f2 double-negative Phox2a cells, which were predominantly located medially, OC2+ Phox2a cells located at intermediate positions, and laterally located Pou2f2+ Phox2a cells (Fig. 5E, F), in line with the expression of OC2 and Pou2f2 in early Phox2a^Deep^ and Phox2a^Sup^ respectively predicted by the adult snRNAseq data (Fig. 5C). We next tested if Pou3f1 is specifically expressed in embryonic late Phox2a^Deep^ cells. Since Phox2a is only expressed in a subpopulation of developing AS neurons ^16^ and due to Phox2a and Pou3f1 antibody incompatibility, we used Lmx1b and Zfhx3 co-expression to label all presumptive AS neurons within the dI5 domain (Fig. 5H). Co-staining at e13.5 confirmed good agreement between Phox2a and Zfhx3 expression in dI5 neurons (Fig. S11A-C). We therefore quantified Pou2f2 and Pou3f1 expression in Lmx1b Zfhx3 double-positive neurons at e13.5. Within the region containing Phox2a^Sup^ neurons migrating into lamina I, 80% AS neurons expressed Pou2f2 and essentially none expressed Pou3f1 (Fig. 5G). Meanwhile, 58% of medial cluster AS neurons containing the late Phox2a^Deep^ neurons expressed Pou3f1 and virtually no Pou2f2 (Fig. 5G). Together, our data reveal a correlation between the OC2 > Pou2f2 > Pou3f1 expression sequence and early AS^Deep^ > AS^Sup^ > late AS^Deep^ birth sequence.

### Pou2f2 and Pou3f1 are required for the normal specification of limb-innervating motor columns

We next tested whether the expression of OPP proteins is required for the diversification of select populations of spinal neurons. Specifically, we asked whether the genetic loss of an OPP protein results in: (1) the impaired specification of the neuronal population normally expressing it, and/or (2) its conversion into the population normally expressing the next or the previous OPP in the temporal sequence. To address this, we converted the conditional *Pou2f2* allele, *Pou2f2^flox^* ^66^, to a constitutive *Pou2f2* loss of function allele (*Pou2f2*^-^) using a germline Cre source. The effects of Cre-mediated excision on Pou2f2 protein levels and localization were further characterized with two different anti-Pou2f2 antisera (Fig. S11D and Experimental Procedures), suggesting *Pou2f2*^-^ is a near-null allele.

We first focused on e11.5 forelimb-innervating LMC MNs where Pou2f2 expression is restricted to the LMCl, while Pou3f1 is enriched in the MMC. Compared to control embryos, *Pou2f2^-/-^*mutants had a significantly reduced number of LMCl neurons and a corresponding increase in the number of LMCm neurons, leaving the total number of LMC neurons unchanged (Fig. 6A, B). We did not observe increased numbers of MMC MNs (Mecom+ or Foxp1-/Isl1/2+) normally expressing the next temporal TF in sequence, Pou3f1 (Fig. 6A, B and S12A, B). Finally, LMCl neurons could still be detected at e15.5 (Fig. S12C), indicating that the effect of Pou2f2 loss on LMCl neurons is not greatly exacerbated at later stages of development. We also assessed the effect of Pou2f2 loss on the development of PGC neurons that express Pou2f2 at thoracic levels. We compared the expression of nNos between e13.5 control embryos and *Pou2f2^-/-^*mutants. In the rodent embryonic spinal cord, postmitotic PGC neurons are first located near somatic MNs in the lateral extreme of the ventral spinal cord, and eventually migrate dorsally, where they initiate nNos expression and some of them migrate towards the ventricular zone, constituting medial and lateral populations ^67^. In *Pou2f2^-/-^*mutants the total number of nNos+ neurons was not significantly changed (Fig. 6D; 27/hemisection v. 24/hemisection; p=0.056), but the number of laterally-located nNos+ PGC neurons was decreased (Fig. 6D; 18/hemisection v. 14/hemisection; p=0.04). Taken together, these data argue that Pou2f2 is required for the normal development of LMCl and PGC MNs, two subsets that express this TF during embryonic development.

**Figure 6:**
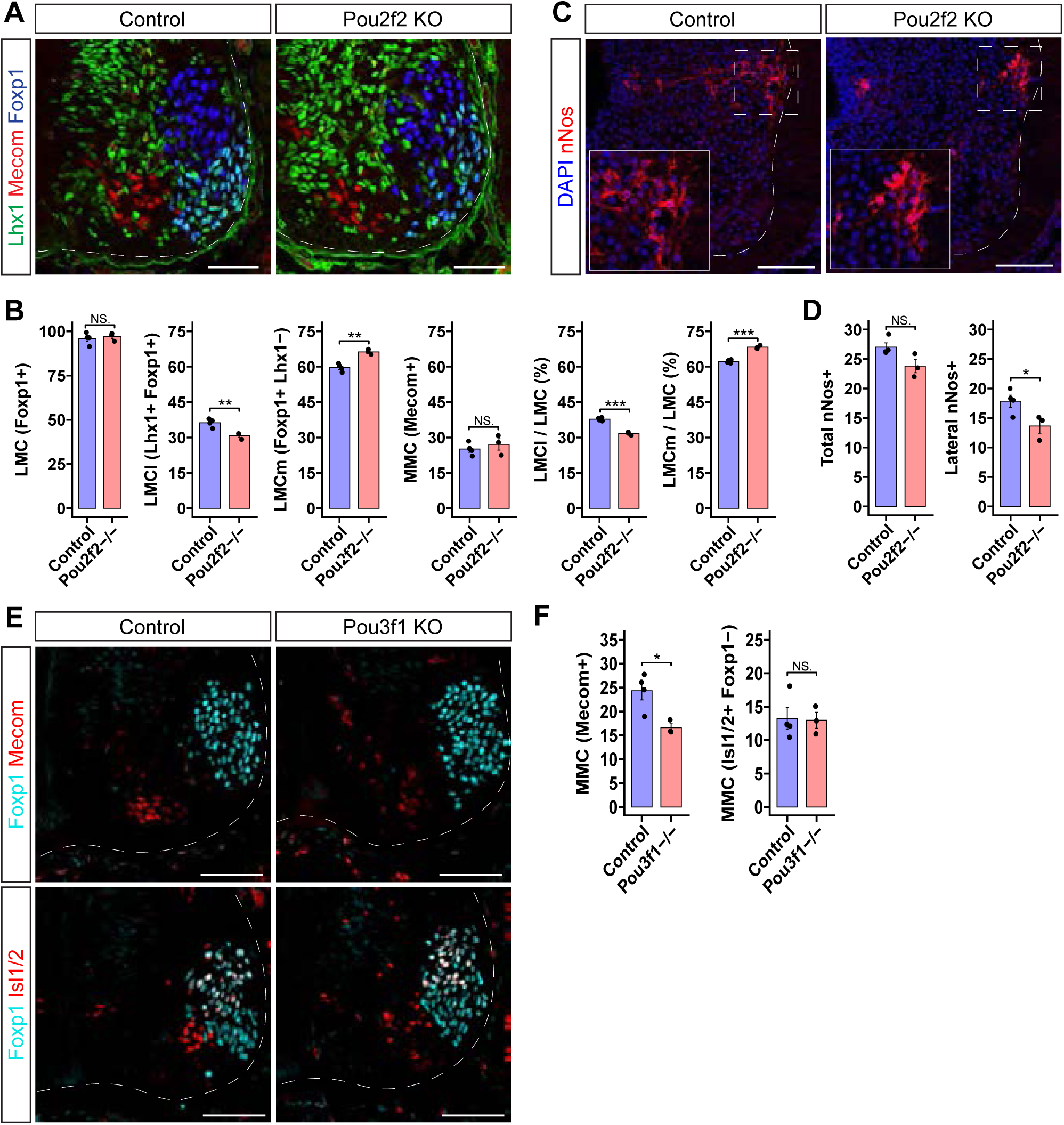
Pou2f2 and Pou3f1 are required for correct motor column specification. (A, B) Reduced number of LMCl neurons and increased number of LMCm neurons in Pou2f2 mutants at e11.5. (A) e11.5 cervical spinal cord sections of control and Pou2f2 mutant embryos stained for the MMC marker Mecom (red), LMC marker Foxp1 (blue) and LMCl marker Lhx1 (green). (B) Number of cells per hemisection for LMCl (i), MMC (ii), LMCm (iii) and LMC (iv) neurons in control and Pou2f2 mutant embryos in cervical spinal cord sections at e11.5. Loss of Pou2f2 causes a significant reduction of LMCl neurons but does not affect the number of MMC, LMCm and LMC neurons. (C, D) Loss of Pou3f1 affects MMC neuron specification. (C) e11.5 cervical spinal cord section stained for the MMC neuron marker Mecom, LMC marker Foxp1 and MN marker Isl1/2. (D) Loss of Pou3f1 causes a significant reduction on the number of Mecom+ MMC neurons (i) but does not affect the number of Isl1/2+ Foxp1-MMC neurons. (E, F) Loss of Pou2f2 affects the specification of thoracic PGC neurons. (E) e13.5 thoracic spinal cord section stained for DAPI and the PGC marker nNOS (red). (F) Loss of Pou2f2 causes a significant reduction of laterally localized nNOS neurons. In B, D, F, Student’s t-test; *p<0.05, **p < 0.01; ***p < 0.001. All data represented as mean ± SEM. Scale bars in A = 50 μm, C and E = 100 μm.

Next, we examined the differentiation of specific spinal motor columns of mouse embryos homozygous for a null/severe loss of function allele of *Pou3f1* (*Pou3f1^-/-^*) ^68^. A comparison of their numbers in *Pou3f1^-/-^* mutants and controls reveals that the loss of Pou3f1 results in significantly fewer Mecom+ MMC neurons (Fig. 6E, F; 24/hemisection v. 17/hemisection; p=0.02) but without affecting the number of Isl1/2+/Foxp1-neurons (Fig. 6E, F; p=0.9), indicating impaired specification of MMC. Furthermore, the loss of Pou3f1 did not result in an increased number of LMCm or LMCl neurons, whose birth precedes MMC neurons (Fig. S12D). Together, our results demonstrate that Pou2f2 and Pou3f1 are essential for the normal development of multiple motor columns.

### Temporal diversification of AS neurons by Pou2f2 and Pou3f1

Next, we examined the development of dorsal horn AS neurons in *Pou2f2^-/-^*mutants. We first focused on superficial AS neurons (AS^Sup^) in which Pou2f2 is highly and specifically expressed (Figs. 7A and S13A). At e15.5 when AS^Sup^ neurons have completed their migration into the superficial laminae and express their terminal differentiation markers, *Pou2f2^-/-^* mutants exhibited a dramatic reduction in the number of superficial Phox2a neurons compared to controls (Fig. 7A, B; p=0.006). This included a reduction in the sparser Phox2a-positive antenna neuron subset of AS^Sup^ which also expressed Pou2f2 (Fig. 7A and S13A,B). To further confirm that this represented a loss of AS^Sup^ neurons rather than a loss of Phox2a expression, we examined a number of other markers of AS^Sup^ neurons. We observed a significant decrease in Zfhx3+ cells in lamina I as well as a similar decrease in superficial cells positive for *Gpr83, Lypd1,* and *Tacr1* mRNA (Fig. S13C, D). Similarly, expression of the TF Pou6f2, which in control animals was expressed in lamina I neurons and a population of Pou2f2+ neurons in the intermediate spinal cord, probably V1 neurons ^69,70^, was lost in the entire spinal cord in *Pou2f2^-/-^* mutants (Fig. S13E), indicating that Pou2f2 is required for the expression of Pou6f2 in all spinal neurons. Meanwhile, the expression of the non-AS specific superficial dorsal horn marker *Reln* was not affected (Fig. S13F).

**Figure 7:**
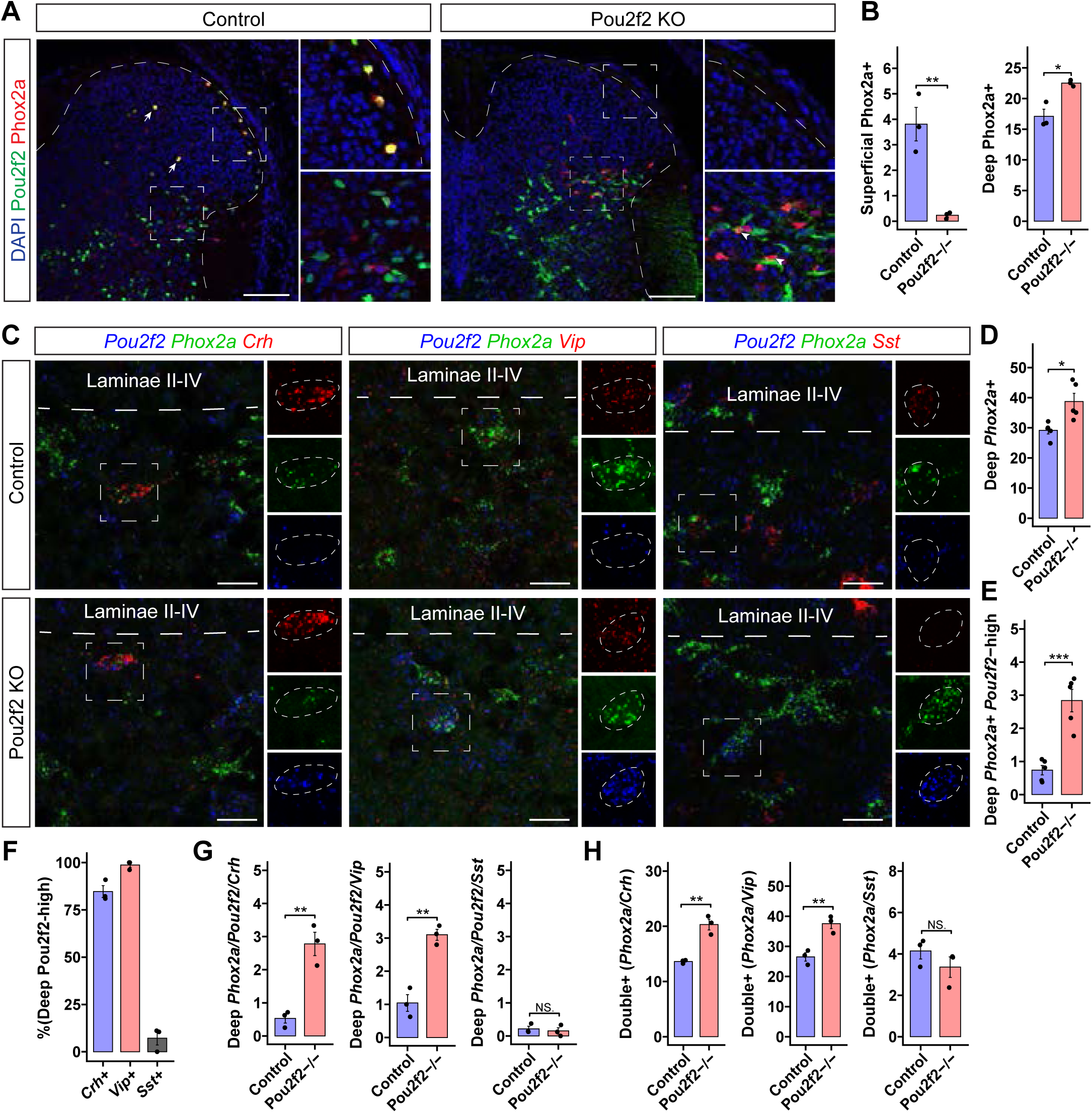
Respecification of Phox2a^Sup^ to Phox2a^DeepLate^ neurons in *Pou2f2* mutants. (A, B) Loss of Phox2a+ lamina I neurons and concomitant increase in lamina V neurons in Pou2f2 mutants (A) e15.5 cervical spinal cord sections of control and Pou2f2 mutants stained for DAPI (blue), Pou2f2 (green) and Phox2a (red). Insets show that loss of Pou2f2 causes near complete absence of Phox2a+ cells in lamina I. Instead Phox2a is detected in neurons in lamina V stained by the truncated cytoplasmic Pou2f2^ΔDNA^ fragment. (B) Quantification of superficial and deep Phox2a+ neurons in control and Pou2f2 mutants at e15.5 (n = 3) (C) RNAscope *in-situ* hybridization for transcripts of *Pou2f2, Phox2a*, and various markers of deep Phox2a AS neurons (*Crh, Vip,* and *Sst*) in controls and Pou2f2 KOs. (D-H) Quantification of number of cells per hemisection within the area of lamina V in C. (D) Increased number of total deep *Phox2a+* neurons in Pou2f2 KOs. (E) Increase in deep *Phox2a+* neurons that highly express the truncated *Pou2f2* transcript in Pou2f2 KOs (these represent putatively converted superficial Phox2a AS neurons). (F) Proportion of deep *Phox2a+/Pou2f2-*high neurons that are positive for each deep Phox2a neuron marker in Pou2f2 KOs. (G) Increase in the number Deep *Phox2a+/Pou2f2-*high neurons positive for *Crh* and *Vip* but not *Sst* in the Pou2f2 KOs. (H) Increase in the total number of *Phox2a+* neurons that are positive for *Crh* and *Vip* but not Sst. In B and D-H Student’s t-test; *p<0.05, **p < 0.01; ***p < 0.001. All data represented as mean ± SEM. Scale bars in A = 100 μm, C = 25 μm.

Concurrent with the loss of AS^Sup^ neurons, we noted an increase in the number of Phox2a neurons located in the deep dorsal horn (Fig. 7A, B; p=0.01), raising the possibility that the loss of Pou2f2 function resulted in the re-specification of AS^Sup^ to AS^Deep^ neurons. To test this possibility, we took advantage of the ability of the antiserum raised against the Pou2f2 N-terminus to recognise Pou2f2^ΔDNA^ produced by the *Pou2f2*^-^ allele (Experimental Procedures). We found a significant increase in Phox2a neurons with cytoplasmic Pou2f2 staining in the deep laminae, in numbers roughly equal to the decrease in superficial Phox2a neurons (Fig. 7B, S13G). Similarly, while there was a slight reduction in the total number of Pou2f2+ Phox2a neurons, potentially due to difficulty detecting the residual cytoplasmic staining, the overall number of Pou2f2+ Phox2a neurons remained similar (Fig. S13H). This led us to assume that Pou2f2^ΔDNA^ can be used to track the development of dorsal horn neurons that were destined to express Pou2f2 in *Pou2f2^-/-^* mutants. To examine their identity, given the absence of AS^Sup^ markers that are not shared by AS^Deep^ neurons, we focused on detection of mRNAs enriched in AS^Deep^ neurons but not in AS^Sup^ neurons (*Crh, Vip, and Sst)* ^16,71^, along with the fact that *Pou2f2* mRNA was detectable in *Pou2f2^-/-^* mutants (Fig. 7C). In line with the Pou2f2^ΔDNA^ localisation data, in *Pou2f2^-/-^* mutants, there was a significant increase in the total number of deep *Phox2a* neurons (Fig. 7D), as well as a significant increase in *Phox2a* expressing neurons co-expressing high levels of *Pou2f2* mRNA in the deep dorsal horn (Fig. 7E). Furthermore, such *Pou2f2*-high *Phox2a*+ neurons expressed *Crh* and *Vip* mRNAs (Fig. 7F, G). Correspondingly, we observed an overall increase in the total number of AS^deep^ neurons that were positive for these markers, as compared to controls (Fig. 7H). Together, these data argue that AS^Sup^ neurons are converted into a AS^Deep^ neurons in *Pou2f2^-/-^* mutants.

Given the apparent re-specification of AS^Sup^ neurons into AS^Deep^ in *Pou2f2^-/-^* mutants, we next asked if Pou3f1 expression was induced in AS neurons in the *Pou2f2^-/-^* mutants. In the e13.5 brachial spinal cord, we observed a significant increase in the proportion of AS neurons that were Pou3f1+ (Fig. 8A, B). We again took advantage of the residual expression of a truncated Pou2f2 protein in the *Pou2f2^-/-^* mutants to assess the proportion of Pou2f2+ neurons that co-express Pou3f1 in the mutants compared to controls. We observed a significant increase in the proportion of double-positive Pou2f2+/Pou3f1+ AS neurons in the mutants (Fig. 8A, B). Together, these data indicate that loss of Pou2f2 results in an activation of Pou3f1 expression in AS neurons, consistent with their conversion to a later-born fate.

**Figure 8:**
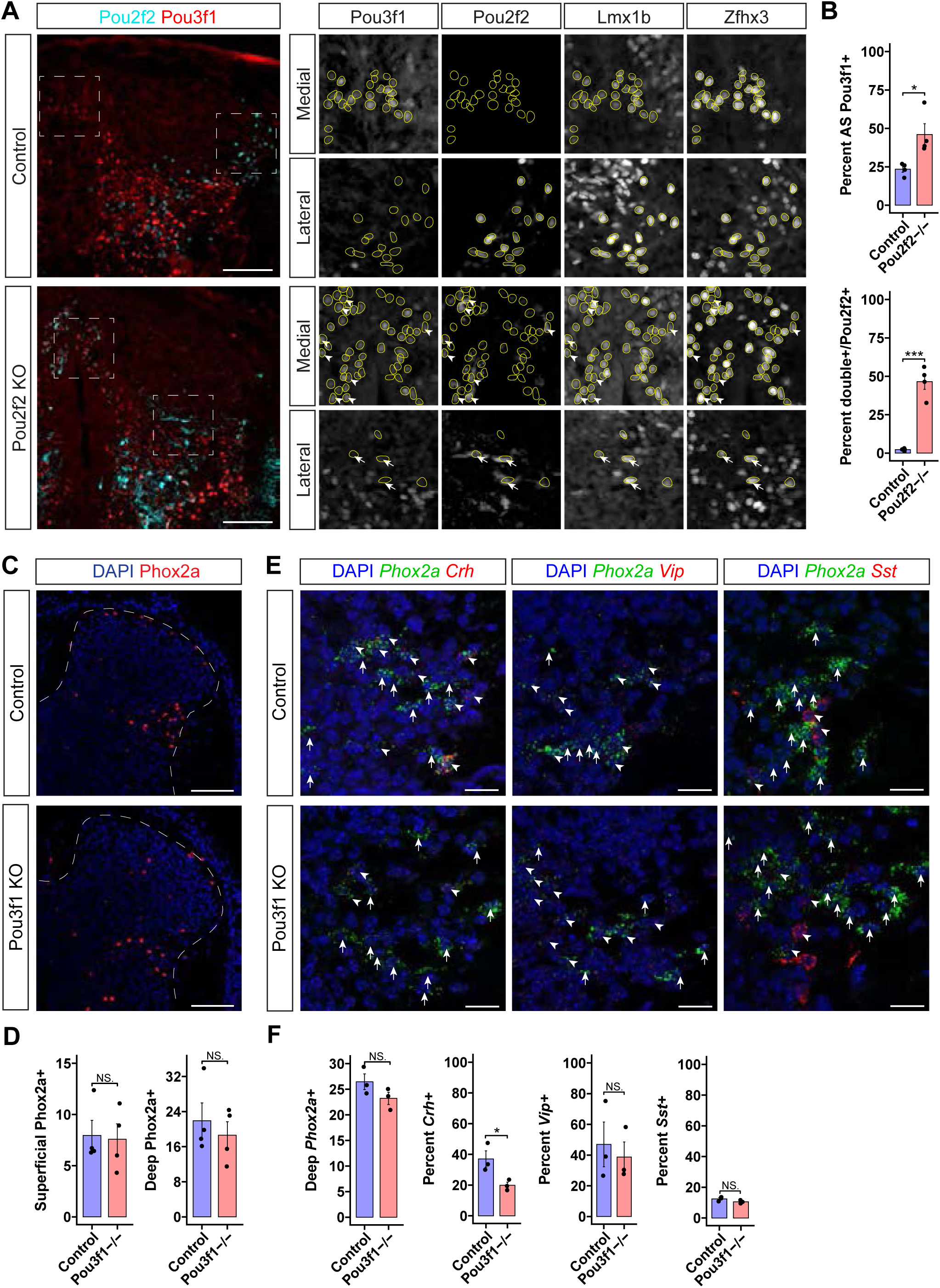
Pou3f1 expression is induced in Pou2f2 KOs and is required for correct specification of deep AS neurons. (A) Staining for Pou2f2 and Pou3f1 in e13.5 brachial sections. While Pou2f2 and Pou3f1 expression is largely non-overlapping in controls, there is significant co-expression of Pou3f1 and the truncated form of Pou2f2 in AS neurons in the Pou2f2 KOs. Yellow circles indicate Lmx1b/Zfhx3 double positive AS neurons. Arrowheads indicate Pou2f2/Pou3f1 co-expressing neurons. Arrows indicate the presence of some Pou2f2 neurons that remain Pou3f1 negative in the Pou2f2 mutants. (B) Quantification of A, showing an increase in the total number of AS neurons (Lmx1b/Zfhx3 double positive) that express Pou3f1 in Pou2f2 KOs compared to controls as well as an increase in the proportion of Pou2f2 positive AS neurons that are also Pou3f1 positive. (C, D) Representative images from e15.5 brachial spinal cords of Phox2a expression in controls and Pou3f1 KOs. (D) Quantification of C showing no change in the number of superficial or deep Phox2a AS neurons. (E, F) RNAscope for *Phox2a* and various deep Phox2a neuron markers (*Crh, Vip,* and *Sst*) in controls and Pou3f1 KOs. (F) Quantification of E showing a significant decrease in the percent of *Phox2a+* neurons that are also positive for *Crh*, but no change in the total number of *Phox2a-*expressing neurons or the percent of them that express *Vip* or *Sst*. In B, D, and F, Student’s t-test; *p<0.05, **p < 0.01; ***p < 0.001. All data represented as mean ± SEM. Scale bars in A, C = 100 μm, D = 25 μm.

Next, we assessed the development of AS neurons in *Pou3f1^-/-^*mutants. We found no change in the number of superficial or deep Phox2a+ neurons (Fig. 8C,D), indicating that in contrast to the *Pou2f2^-/-^* mutants, loss of Pou3f1 did not respecify deep Phox2a+ neurons to the complementary (i.e. superficial) population of AS neurons. This is further supported by the lack of any increase in Pou2f2 expression within Phox2a+ neurons (Fig. S14A, B). The spatial distribution of deep Phox2a+ neurons was similarly unaffected (Fig. S14C). Finally, the proportion of Phox2a+ neurons that were Zfhx3+ was not changed, indicating that their identity as projection neurons was unaffected (Fig. S14A, B). We next assessed the same panel of mRNAs enriched in deep Phox2a neurons as before (*Crh, Vip, and Sst*). We found a significant decrease in the proportion and total number of deep *Phox2a*+ neurons that expressed *Crh* mRNA, while the expression of *Vip* and *Sst* was unaffected (Fig. 8E, F and S14D). Together, these data indicate that Pou2f2 and Pou3f1 are required for the correct specification of their associated subsets of AS neurons, with loss of Pou2f2 being sufficient to drive AS^Sup^ neurons towards a AS^Deep^ fate, possibly through the activation of Pou3f1, the next TF in the OPP temporal sequence.

## Discussion

The generation of neuronal diversity is a fundamental unanswered question of developmental neurobiology. The concept of spatially segregated cardinal domains in the spinal cord giving rise to neurons of related function has withstood many functional validation experiments. Another mechanism of neuronal diversification takes advantage of the chronological order of neuronal birth: different neuronal types emerge in a stereotypical sequence, driven by a succession of transcription factors. Our experiments provide evidence that the archetypal sequence of temporal TFs, OC2 > Pou2f2 > Pou3f1, matches the distinct identities of sequentially generated neuron types within the cardinal domains containing MNs and AS neurons. By studying mice lacking Pou2f2 and Pou3f1 function, we reveal that these TFs are involved in the differentiation of distinct neuronal subtype identities within two cardinal domains. Here, we discuss our findings in terms of the similarities and differences in the strategies of temporal patterning of motor neurons, motor-related interneurons and anterolateral system neurons, and the interplay between spatial and horologic diversification of spinal neurons.

### A temporal logic of transcriptional identity of motor-related cardinal groups in the spinal cord

Our observations provide evidence that Pou2f2 and Pou3f1 are required for normal MN development and that the OC2 > Pou2f2 > Pou3f1 expression sequence in MNs correlates with the LMCm > LMCl > MMC chronology of MN generation. Neither the loss of Pou2f2 nor that of Pou3f1 results in significant changes in the total number of MNs, indicating that they are not required for the generation or consolidation of the broad MN identity. Rather, in mice lacking Pou2f2 function, some LMCl MNs appear to be respecified into the antecedent columnar type in the sequence, the LMCm MNs. At non-limb levels, the loss of Pou2f2 also affects PGC specification, although it is less clear whether these MNs are respecified into a particular columnar type. The loss of Pou3f1 disrupts MMC MN differentiation, but we did not find evidence of their conversion into another MN columnar type as defined by standard marker expression. These data contrast a previous study in which the loss of OC2 results in a re-specification of an earlier motor column identity into the following one in the sequence, i.e., LMCm into LMCl MNs ^40^. One potential explanation is that the increase in LMCm neuron numbers in Pou2f2 mutants may result from a subpopulation of LMCl MNs that co-expresses Pou2f2 and OC2, taking on an LMCm identity (OC2+) in the absence of Pou2f2. However, more analysis will be needed to fully appreciate the specific role of Pou2f2 (and Pou3f1) in MN development, especially in the context of a hierarchy of other potentially redundant temporal TFs.

Foxp1 is required for the specification of the LMC and PGC, both early-generated MN subtypes at brachial and thoracic levels respectively ^8,10,72^. Recent observations also argue that Foxp1 is involved in the specification of early neuronal identities in the cortex and retina ^73,74^. Thus, Foxp1 appears to represent a temporal TF that specifies early identities in the MN lineage at different axial identities, i.e., instructed by the Hox TF context ^8,10^. This interpretation also provides an explanation for the puzzling observation that LMC and PGC neurons are both transformed into seemingly unrelated HMC neurons in Foxp1 mutants – the only early Foxp1-negative MN subtype. Moreover, the interplay between OPP and Foxp1/Lhx3 TF programs likely involves cross-repressing interactions, as documented between Foxp1 and Pou3f1 ^10,75^. This cross-repression may underlie the observed switch from Pou2f2 to Pou3f1 expression coinciding with the LMC (Foxp1+) to MMC (Mecom+) identity transition, and why in contrast to AS neurons, loss of Pou2f2 does not cause precocious induction of Pou3f1 in MNs. Therefore, we suggest a potentially concurrent role for Foxp1 at a similar temporal window as OC2 and Pou2f2 at brachial levels and as Pou2f2 at thoracic levels, where this temporal overlap, in conjunction with the combinatorial activity of Hox TFs and cross-repressive interactions among TFs, orchestrates the acquisition of distinct motor column identities along the axial spinal cord.

Our experiments reveal an OC2 > Pou2f2 > Pou3f1 chronology of expression consistent between the different cardinal domains. This sequence implies an LMCm > LMCl > MMC birth order, reconciling previous LMC MN birth-dating observations with a more speculative MMC birth time ^34,35,40^. Relying solely on Isl1 and Lhx3 expression as a molecular signature of MMC neurons suggests an MMC > LMC birth order since all newly born MNs co-express Isl1+ Lhx3+ ^11,60^. However, this approach provides an incomplete picture of the progression of MMC specification. Considering the expression of Mecom, a marker of maturing MMC neurons in the migratory stream linking the pMN domain with the MMC column, together with the expression of Pou3f1 revealed by our experiments, argues for MMC specification following that of LMC neurons.

The OPP temporal TF sequence also parcellates motor-related interneurons into molecularly and functionally distinct subtypes. For example, in V1 neurons, Onecut TFs are required for the specification of Renshaw cells ^32^, and the expression of Pou6f2, which defines a clade of V1 neurons ^69^, is lost in the spinal cord of Pou2f2 mutants, as per our analyses (Fig. S13, data not shown). This situation is similar to V3 neurons, where OC2 and Pou2f2 define molecularly, spatially and anatomically distinct subsets of V3 neurons, V3_D_ and V3_V_ ^63^. The diversification of V3 neurons appears to follow the OC2 > Pou2f2 > Pou3f1 sequence since the switch from Pou2f2 to Pou3f1 expression observed by us coincides with the switch in expression from the V3_D_ marker Lhx1 to the V3_V_ marker Olig3, aligning with the known V3_D_ > V3_V_ chronology of birth ^30^.

### Temporal patterning of AS neuron subtypes

Neurons that express the transcription factor Phox2a during development constitute a substantial fraction of the AS, essential for the relay of somatosensory information from the spinal cord to the brain ^16^. Based on their highly diverse migration patterns, final location and molecular profile, developing Phox2a AS neurons expressing the transcription factor Phox2a can be partitioned into at least five distinct subsets, spanning the superficial and deeper layers of the dorsal horn. Their birthdating suggests the following sequence of ontogenesis: early AS^Deep^ > AS^Sup^ > late AS^Deep^, and our observations show it as corresponding to the OC2 > Pou2f2 > Pou3f1 expression sequence. This evidence is bolstered by the fact that OC2 and Pou2f2 expression persists, respectively, in the early-generated AS^Deep^ neurons and AS^Sup^ neurons ^16,65^. Our experiments also show that Pou2f2 is expressed in all AS neurons in the superficial laminae, including non-Phox2a projection neurons, likely the Tac1-expressing AS^Sup^ population that is complementary to Phox2a-expressing neurons; thus, the chronology of OPP expression may direct the development of all AS neurons.

We tested the functional significance of the OPP expression chronology by studying mice with a loss of function of Pou2f2 and Pou3f1 and found that both are essential for the development of the AS subsets that expresses them. The loss of Pou2f2 resulted in a re-specification of AS^Sup^ to AS^Deep^ late fate since neurons fated to express Pou2f2, (1) do not enter the superficial layers of the dorsal horn, (2) express neuropeptide-encoding mRNAs characteristic of AS^Deep^ late neurons and (3) maintain their projection neuron fate by expressing Zfhx3. Interestingly, the AS^Deep^ late fate is the next one in the AS developmental sequence, echoing the findings in OC2 mutant mice, where earlier born MN take on the next fate in MN developmental sequence ^40^. This observation also suggests that Pou2f2 directly or indirectly represses the deep-late transcriptional program, and its absence causes the precocious induction of expression of Pou3f1, although the extent to which this effect contributes to the observed AS^Sup^ → AS^Deep^ late fate switch is unclear. The loss of Pou2f2 affects the development of all AS^Sup^ neurons, including those that do not express Phox2a; hence, this manipulation may provide a way to explore the relative contribution of AS^Sup^ and AS^Deep^ neurons to somatosensation.

The loss of Pou3f1 reveals that it is required for the normal expression of at least one of the neuropeptide mRNAs typical of AS^Deep^ late neurons. Also, in the absence of Pou3f1, AS^Deep^ late neurons do not take on the AS^Sup^ fate as there are no supernumerary AS neurons in the superficial dorsal horn, or increased Pou2f2 expression. What then is the fate of AS^Deep^ late neurons lacking Pou3f1? Within the broader context of dI5 cardinal domain Lmx1b+ neurons, Phox2a neurons form the projection neuron group within the projection neurons > interneurons temporal scheme ^37^. However, in the absence of Pou3f1, AS^Deep^ apparently maintain the expression of Zfhx3, a projection neuron marker, and the expression of Phox2a, required for AS neuron connectivity to the brain, thus it is unlikely that they are respecified to become interneurons.

Our data establish temporal identity as an organizing principle that not only segregates long-range neurons from local interneurons but also parcellates long-range projection neurons into distinct subsets. Given the conservation of the OPP TF sequence in multiple classes of spinal cord neurons, it is likely that these TFs are also involved in the specification of distinct projection neuron subsets within these classes. In support of this hypothesis, recent mapping of the projection patterns of OC2+ and Pou2f2+ V3 neuron subsets revealed distinct settling and projection patterns of these neurons ^63^. The expression of OPP TFs is conserved in large swathes of the developing nervous system ^39^, and there is at least one study providing functional significance of this: Pou3f1 is required for the specification of contralaterally projecting retinal ganglion cells, a major projection neuron population ^76^. Thus, the OPP TF sequence directing the specification long-range projection neuron subtypes could be a general diversification mechanism in the developing nervous system.

### Combinatorial superposition of spatial and temporal TF codes: a robust neuronal diversification strategy

Our expression profiling and functional experiments provide evidence that the OPP temporal TF sequence is deployed within multiple cardinal domains resulting in a spatio-temporal intersection of two molecular programs of neuronal diversification. Comparing these findings with the classical observations in *Drosophila* offers a glimpse into the developmental logic underlying this intersection. The spatio-temporal diversification strategy in *Drosophila* involves specifying distinct neuroblast spatial identities and their subsequent patterning by common sequences of temporal TFs ^21,77,78^. While we observe some expression of OPP TFs in spinal progenitors (Fig. S2A,B), they are predominantly expressed in post-mitotic neurons with multiple OPP TFs being coexpressed and eventually resolving to single OPP TFs. This suggests that early postmitotic neurons are competent to express all three TFs and to acquire the associated neuronal identities. Beyond these general inferences, the specific time window of OPP TF requirement in spinal neurons remains speculative; however, it may be mechanistically advantageous to segregate the timing of cardinal type specification at the level of progenitors, from the temporal diversification of post-mitotic neurons. Another important difference between *Drosophila* and spinal cord strategies is that in *Drosophila*, temporal TFs function by activating the next one in the series while repressing the previous one. In contrast, in the spinal cord, the opposite logic is apparent: early TFs repress subsequent temporal identities and their loss results in the re-specification to later temporal identities, as implied by the loss of OC TFs resulting in a LMCm → LMCl conversion ^40^, and the loss of Pou2f2 causing a AS^Sup^ → AS^Deep^ late identity. Also, our observations argue against the loss of an OPP TF resulting in the promotion of an earlier neuronal identity. For example, the loss of Pou3f1 does not promote earlier LMC or AS identities, and the loss of Pou2f2 only causes a limited increase in the number of LMCm neurons, suggesting that OPP TFs do not restrict the temporal windows during which earlier neuronal identities are specified. Alternatively, each phase may be determined by multiple TFs and thus removal of individual TFs may not be sufficient to expand the temporal windows of earlier identities. Finally, while many characteristics of the OPP TFs are consistent with temporal TFs in *Drosophila*, other characteristics are more reminiscent of terminal selector genes ^79^, indicating that OPP TFs may reside at an intermediate level of the fate specification hierarchy between temporal patterning of progenitors and terminal fate selection. Thus, while the general developmental logic of spatio-temporal intersection appears to apply to insect and mammalian spinal cord neurons, the progression of the temporal TF series appears to be dictated by different molecular mechanisms to be unravelled by future studies.

## Experimental Procedures

### Key resources table

INSERT HERE – separate file in the manuscript folder

#### Animal welfare

Animal experiments in the Kania lab were carried out in accordance with the Canadian Council on Animal Care guidelines and approved by the IRCM Animal Care Committee. Animal experiments in the Sagner lab were performed in accordance with German and European animal welfare laws.

#### Generation and characterization of the *Pou2f2^-^* allele

*Pou2f2^flox^* has LoxP sites surrounding *Pou2f2* exons 8-11 that encode the DNA binding domain ^66^. This conditional allele was converted to a constitutive loss of function allele using the germline cre EIIa-Cre ^80^. The effect of Cre-mediated excision of Pou2f2 protein expression was validated using 2 antibodies (Fig. S11D). One raised against the N-terminal domain preceding the DNA domain revealed a relocalisation of the truncated Pou2f2 (Pou2f2^ΔDNA^) from the nucleus to the cytoplasm, consistent with the generation of a truncated protein lacking the POU DNA binding domains and requirement of these domains for nuclear localization ^66^. This was observed in dorsal and ventral neurons, but not in MNs, possibly due to the generally lower Pou2f2 expression in MNs (Figs. 4L-N and top row in S11D), and the large volume of their cytoplasm diluting the Pou2f2^ΔDNA^ beyond the limit of detection. The second antiserum did not detect Pou2f2^ΔDNA^, suggesting that it was generated against a C-terminal domain of Pou2f2 (Fig. S11D; see Materials and Methods for details). Together, these argue that the excision of exons 8-11 generated a near-null allele of *Pou2f2* (*Pou2f2*^-^).

#### Genotyping

Genotyping by PCR for the Pou2f2^fl^ allele was performed using the following primers: forward: 5’-CCATCTGTAGTCCAGGCATC-3’ and reverse: 5’-GACATTCCCCTCCATTGAAC-3’, producing a wild-type amplicon of 170 bp and a mutant amplicon of 310 bp. Genotyping of the Pou2f2^-^ allele was performed using the same forward primer as the Pou2f2^fl^ allele, with an additional reverse primer: 5’-ATGTGCCGGTCACAACTCTT-3’, producing a mutant (KO) amplicon of 434 bp that was distinguishable from the flox amplicon. Genotyping for the Pou3f1^-^ allele was performed using the following primers: Pou3f1 WT allele: forward: 5’-CAAGCGCAAGAAGCGCACGTCCATC-3’, reverse: 5’-GTTGCAGAACCAGACACGCACCACC-3’, additional Pou3f1 mutant allele Forward primer: 5’-CTTCCTCGTGCTTTACGGTATCGC-3’. These produced a WT band at 154bp and mutant Band at 320bp. An additional independent primer pair for the Pou3f1 WT allele was also used (Forward: 5’-AAGCAGTTCAAGCAACGACG-3’, Reverse: 5’-TAACTGCGCGCCGGA-3’) with the addition of Q-solution (Qiagen, Cat#210220).

#### Tissue collection

On the appropriate embryonic day to each experiment, the pregnant female mouse was anesthetized with 0.3 ml of 10 mg/ml Ketamine, 1 mg/ml Xylazine in 0.9 % saline prior to cervical dislocation. Embryos were dissected in ice cold 1x Phosphate-buffered saline (1x PBS – 0.1 M Na_2_HPO_4_, 0.1 M NaH_2_PO_4_, 0.15 M NaCl) (pH 7.4), tissue samples were taken for genotyping, and then embryos were fixed for 3 hours in 4 % PFA in 1x PBS at 4 °C while mechanically shaken. Post-fixation, embryos were washed in 1x PBS followed by a two-day incubation in 30% sucrose in 1x PBS or over-night incubation in 15% Sucrose in 0.12 M PB for cryoprotection. After embryos had sunk in the sucrose solution, they were immersed in OCT compound (Sakura Finetek #: 4583), frozen in blocks of OCT at −80 °C or dissected in 15% Sucrose in 0.12 M PB solution, incubated in gelatine solution, deep-frozen in isopentane, and stored at −70 - −80 °C. 12 – 14 µm cryosections were cut at −20 °C. Tissue sections were subsequently stored at −70 - −80 °C.

#### Immunohistochemistry

Slides were warmed to room temperature and washed 3 times with 1x PBS, 5 minutes each, and incubated in a blocking solution (0.1% Triton X-100 in 1x PBS (0.1% PBS-T) and 5% heat-inactivated horse serum (HIHS) or 1% BSA) for 30 minutes to 1 hour. Sections were then incubated with primary antibody solution (primary antibodies diluted in blocking solution) overnight at 4 °C. On the following day, sections were washed 3 times with 1x PBS for 5 minutes or 3 times with PBS-T for 30 minutes and incubated with secondary antibody solution at room temperature for 1-2 hours. Following the incubation, sections were washed 3 times with 1x PBS for 5 minutes or 3 times with PBS-T for 30 minutes and coverslipped using a Mowiol solution (10% Mowiol, 25% glycerol; Sigma). Slides were allowed to dry and stored at 4 °C in the dark prior to imaging. For a list of antibodies used in this study, please see the Key Resources table.

#### RNAscope In situ-hybridization (ISH)

ISH was performed using RNA Scope Multiplex Fluorescent v2 kits according to manufacturer’s (ACDBio) instructions. Probes used are as follows: Mm-Lypd1-C1 (ACD Bio: #318361), Mm-Tacr1-C1 (ACD Bio: #410351), Mm-Gpr83-C1 (ACD Bio: #317431), Mm-Crh-C1 (ACD Bio: #316091), Mm-Vip-C1 (ACD Bio: #415961), Mm-Sst-C1 (ACD Bio: #404631) Mm-Pou2f2-O1-C2 (ACD bio: #1170721-C2) Mm-Phox2a-C3 (ACD Bio: #520371-C3), and Mm-Reln-C3 (ACD Bio: #405981-C3).

#### EdU and BrdU birth-dating

For EdU labeling, mice were intraperitoneally injected with 3 μl/g body weight with EdU diluted in PBS at a concentration of 10 mg / ml at the indicated stages. EdU was detected using Alexa647 Click-iT EdU Imaging Kit (Invitrogen C10340) according to the manufacturer’s specifications.

For BrdU labelling, mice were intraperitoneally injected with 100 μg BrdU (Sigma, #B5002) diluted in PBS. Slides were warmed to room temperature and washed 2 times with 1x PBS-NH4Cl (50mM), 10 minutes each. To perform heat induced antigen retrieval, slides were immersed in a staining dish containing 10 mM Citrate Buffer (0.1M Citric Acid, 0.1M Sodium Citrate, pH 6.0), heated with a Pascal Pressure chamber for 45 minutes and cooled down to room temperature. 5 minutes wash with Milli-Q water was performed, followed by 2 times washes with 1x PBS. Sections were permeabilized in 0.1% PBS-T for 15 minutes at room temperature. Blocking and antibody incubation were performed as previously described. At least 3 sections from different animals were analyzed for each time point.

#### Microscopy and imaging

Immunohistochemistry images were acquired using Leica DM6 or SP8 confocal microscope or Zeiss Axio Observer microscope equipped with an Apotome.2 structured illumination module and a 20× air objective (NA = 0.8). RNAscope images for quantification were acquired on the Leica SP8 confocal microscope at 40X magnification to be able to resolve individual puncta.

#### Image quantification and statistical analysis

All cell counts were performed with Fiji/ ImageJ and recorded to Microsoft Excel. Data was then analyzed further and plotted using R (version 4.0.5) and the R packaged ggplot2 (version 3.4.2). For quantification of RNAscope images, a cell was considered positive for a given mRNA if it contained at least 5 puncta that could be positively assigned to that cell and “high” for a given mRNA if it contained at least 20. For some mRNAs which were less highly expressed overall (Gpr83 and Tacr1) a threshold of 3 puncta was used for a cell to be considered “positive”. For analysing the 2D distribution of deep Phox2a+ neurons in the Pou3f1 mutants, the X and Y coordinates from the ImageJ cell counter plugin were exported to an XML file. Coordinates were then imported into R and normalised to the location of markers placed at the dorsal, ventral, left lateral, and right lateral extremes of the spinal cord. For datapoints on the left hemisection, their coordinates were flipped across the midline to align with those from the right hemisection. Density plots were generated using the geom_density_2d function of ggplot2 with bins=5.

Nuclei were segmented using StarDist and subsequently manually corrected in Fiji. Intensities of individual channels were measured using the Multi Measure function of Fiji’s ROI Manager and exported as csv or Microsoft Excel files. Data analysis and plotting were performed in R (https://www.r-project.org). For each section, intensities in nuclei were first normalized between 0 and 1. Intensity thresholds corresponding to 0 and 1 were determined by discarding intensities of a fraction of the brightest and dimmest objects, typically 0.1 %. Objects were counted as positive for expression of temporal TFs if their normalized intensities exceeded a normalized intensity of 0.2, for Neurog2, lacZ and EdU a normalized intensity of 0.3. For assigning motor column identities, normalized intensity thresholds of 0.15 for Isl1 and 0.125 for Foxp1 were used. Plots were generated using ggplot2 or Prism software.

Test methods and p values were described in figure legends, with p value 0.05 as a significance threshold.

#### Image quantification and statistical analysis

scRNAseq analysis was performed in R-Studio using R v3.5.2 and later. Scripts describing the scRNAseq analysis performed in this manuscript are available as Supplemental Experimental Procedures.

##### General OPP expression dynamics and identification of Pou3f1 as temporal TF

scRNAseq data from e9.5 to e13.5 spinal cord neural progenitors including subtype annotations were obtained from Delile et al. 2019 (E-MTAB-732). The annotation of a subset of neurons, formerly annotated as dI4, was altered to dI6 based on marker gene expression and overlap with the few annotated dI6 neurons in the initial dataset. Differential gene expression between neurons from different embryonic days was performed using Seurat’s "FindAllMarkers” function. To count the number of DV classes in which the identified TFs are differentially expressed, differential gene expression analysis between neurons from different embryonic days was performed on neurons in each class using the “FindAllMarkers” function with settings min.pct = 0.25 and logfc.threshold = 0.25.

##### OPP expression in motor and V3 neurons

For analysis of MNs at e10.5 and e11.5, the scRNAseq dataset of Delile et al. 2019 was used. For analysis of e12.5 MNs and V3 neurons that dataset from Wang et al. 2022 was used (GSM4734731). For the Delile dataset, neurons of the respective stages and identities were identified based on the provided annotations. For the Wang dataset, motor and V3 neurons were identified based on cell clustering and marker gene expression. UMAP-based dimensionality reduction was performed in Seurat using the first 30 principle components. Cell clusters were identified using Seurat’s FindNeighbors and FindClusters functions. Expression of established marker genes was subsequently used to assign cell identities to clusters. Dot plots were generated using Seurat’s DotPlot function. The Monocle3 software package was used for pseudotime reconstruction of V3 neurons.

##### OPP expression in adult AS neurons

Cell type annotations and scRNAseq data of Bell et al. 2024, provided via the GEO repository (GSE240528), were directly loaded. OPP expression levels in the annotated ALS1-5 clusters was plotted using Seurat’s DotPlot function.

## Supporting information

Key resources table

## Data Availability

Further information and requests for resources and reagents should be directed to and will be fulfilled by the lead contacts, Artur Kania and Andreas Sagner. Any additional information required to reanalyze the data reported in this work is available from the lead contacts upon request.

## Acknowledgments

We thank Michel Cayouette, Michael Wegner, R. Brian Roome and James Briscoe for comments on the manuscript, Dominic Fillion for assistance with microscopy, Meirong Liang for technical support, and Michel Cayouette, Christine Jolicoeur, Frederic Clotman, Michael Wegner, Elisabeth Sock, Chichung Lie and James Briscoe for sharing mouse lines and antibodies. We are also grateful to all members of the Sagner and Kania labs, past and present, for help, advice and critical feedback. This work was supported by grants from the German Research Foundation (DFG; project number 455354162) and the Interdisciplinary Centre for Clinical Research (IZKF – E36) to AS, and project grants from the Canadian Institutes of Health Research (PJT-162225, MOP-77556, PJT-153053, and PJT-159839) to AK. KTS is a recipient of a CIHR doctoral scholarship (FRN:494078). AK holds the Doggone Foundation Chair of Excellence in Pain.

## Author contribution

All authors: Conceptualisation, Writing – Editing and Review.

LCS: Methodology, Investigation, Data Curation, Writing – Original Draft, Visualisation, Resources.

KTS: Methodology, Investigation, Data Curation, Visualisation, Writing – Original Draft.

JS: Methodology, Investigation, Data Curation, Visualisation.

AK: Supervision, Funding Acquisition.

AS: Writing – Original Draft, Supervision, Project Administration, Funding Acquisition.

## Ethics

All vertebrate animal experiments were carried out in accordance with the Canadian Council on Animal Care guidelines and approved by the IRCM Animal Care Committee (Protocol 2019-09 AK and 2021-12 AK).

## Supplemental Figure Legends

**Figure S1:**
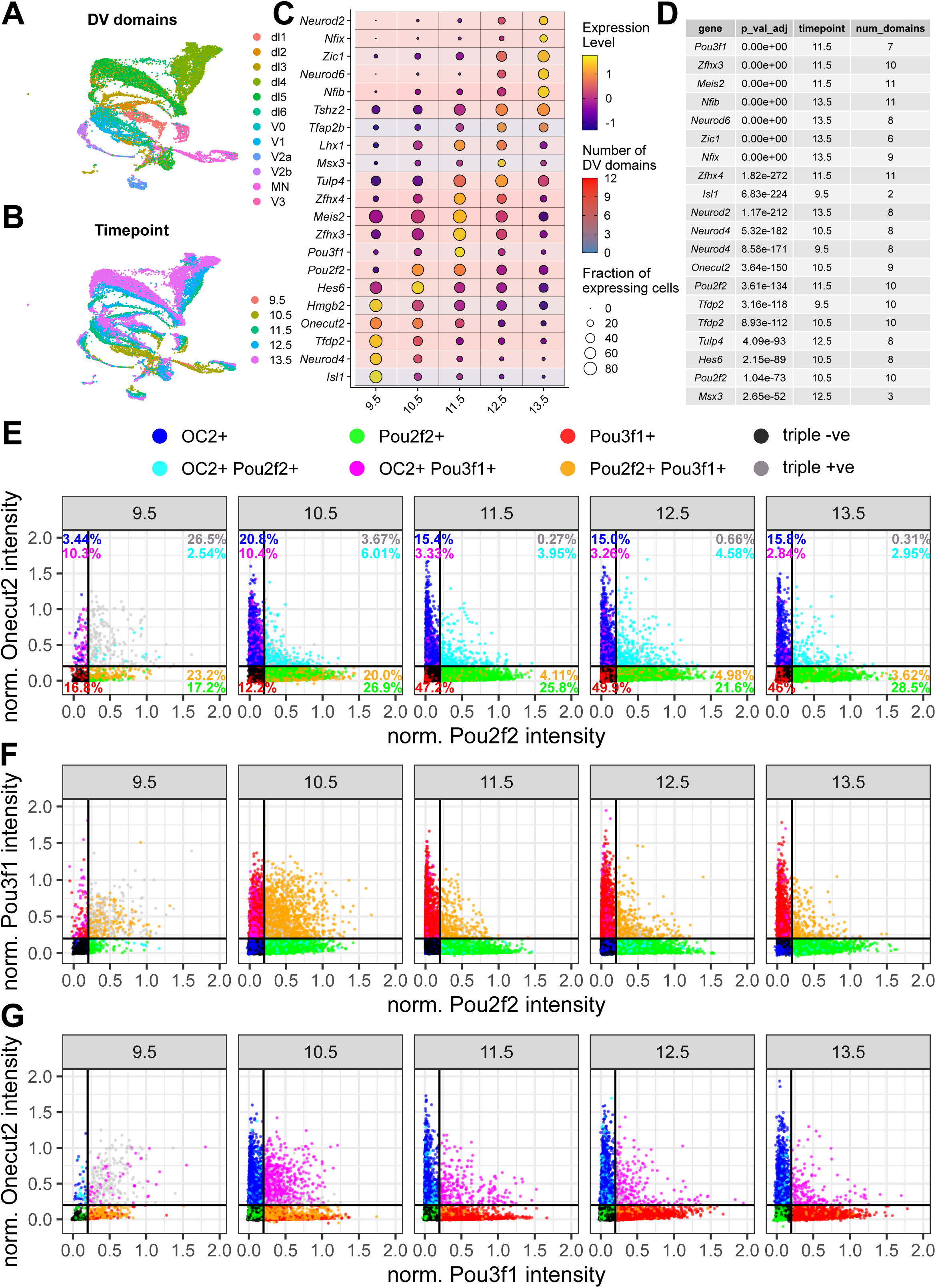
Expression dynamics of OPP TFs during spinal cord development (Related to Figure 1) (A - D) Chronological expression of Pou3f1 in scRNAseq data from Delile et al. 2019. (A, B) UMAP plots showing DV class and developmental stage of neurons. (C) Expression dynamics of TFs differentially expressed in neurons sampled between e9.5 and e13.5. Color and size of the circles denote scaled expression levels and fraction of expressing cells. The color of the background indicates the number of individual DV classes in which the TF was differentially expressed between neurons from different embryonic days. (D) Results of the differential gene expression analysis. For each TF, adjusted p-value, embryonic day at which the TF was annotated as differentially expressed, and number of DV domains in which the TF was differentially expressed is provided. (E - G) Intensity distributions of OPP TFs from e9.5 to e13.5.

**Figure S2:**
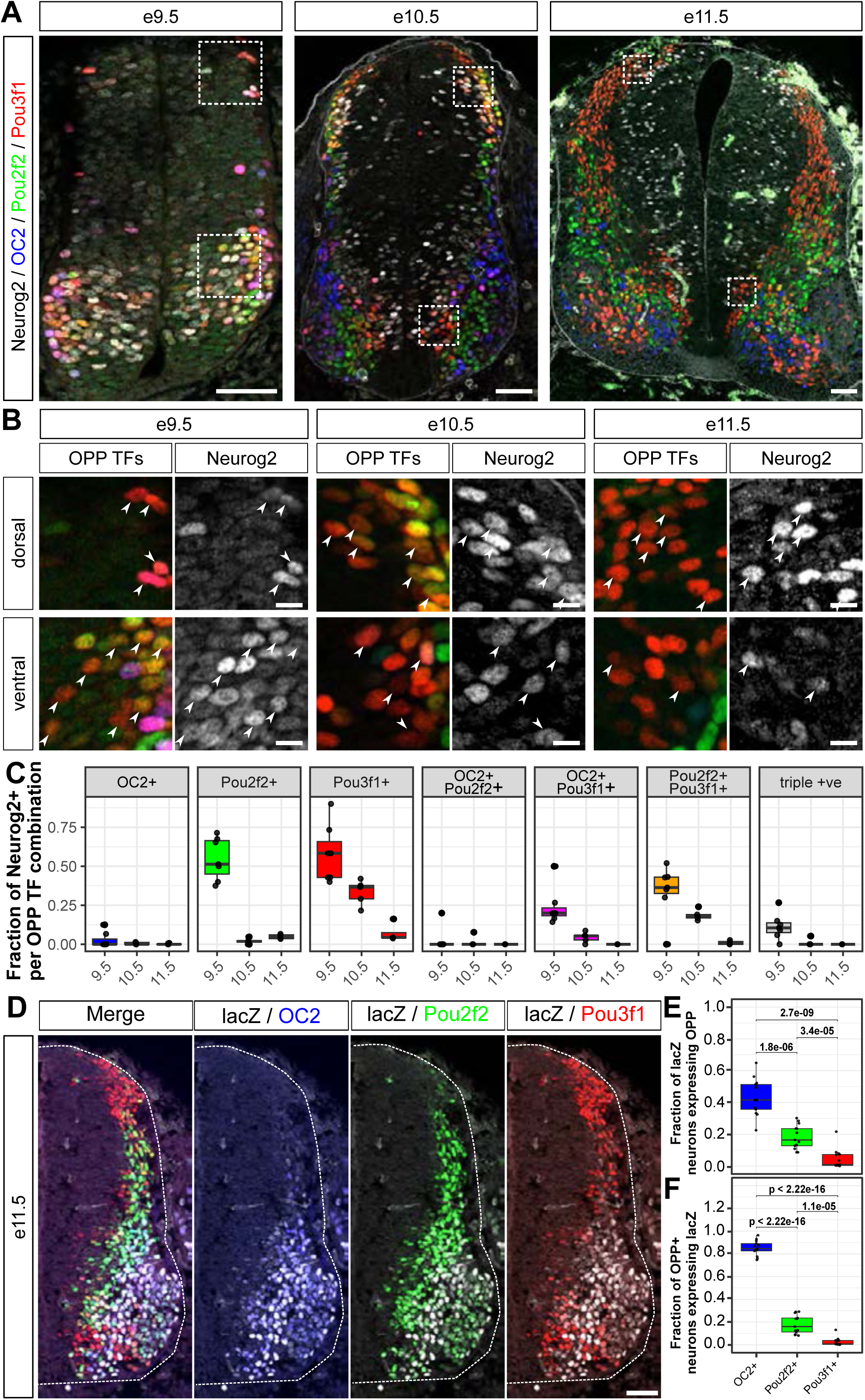
Transient co-expression of OPP TFs shortly after neurogenesis (Related to Figure 1) (A) e9.5 (left), e10.5 (middle) and e11.5 (right) spinal cord sections co-stained for OPP TFs and Neurog2. (B) Higher magnification views of the regions indicated in (A). White arrowheads indicate cells co-expressing Pou3f1 or multiple OPP TFs and Neurog2. (C) Quantification of the fraction of cells per OPP TFs combination that co-express Neurog2. Scale bars in A, D = 50 µm; in B = 10 µm

**Figure S3:**
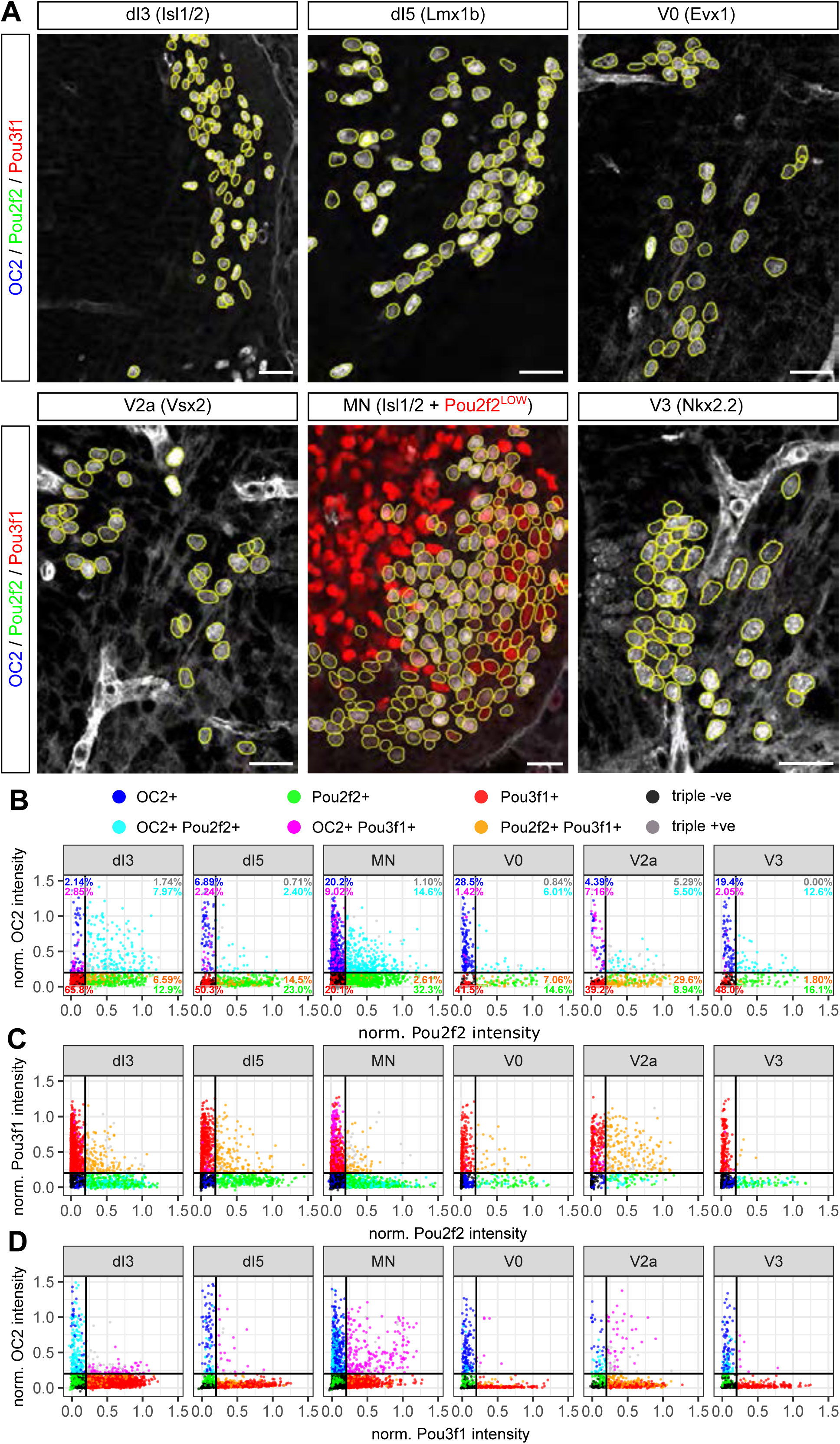
OPP TFs parcellate spinal neuronal classes into distinct subtype identities (Related to Figure 2) (A) Segmentation of neurons with specific DV identities (yellow outlines) were obtained from co-staining with the indicated DV markers (same sections as Fig. 2A). (B) Intensity distributions of OPP TFs in different DV classes of neurons at e11.5. Scale bars in A = 100 µm

**Figure S4:**
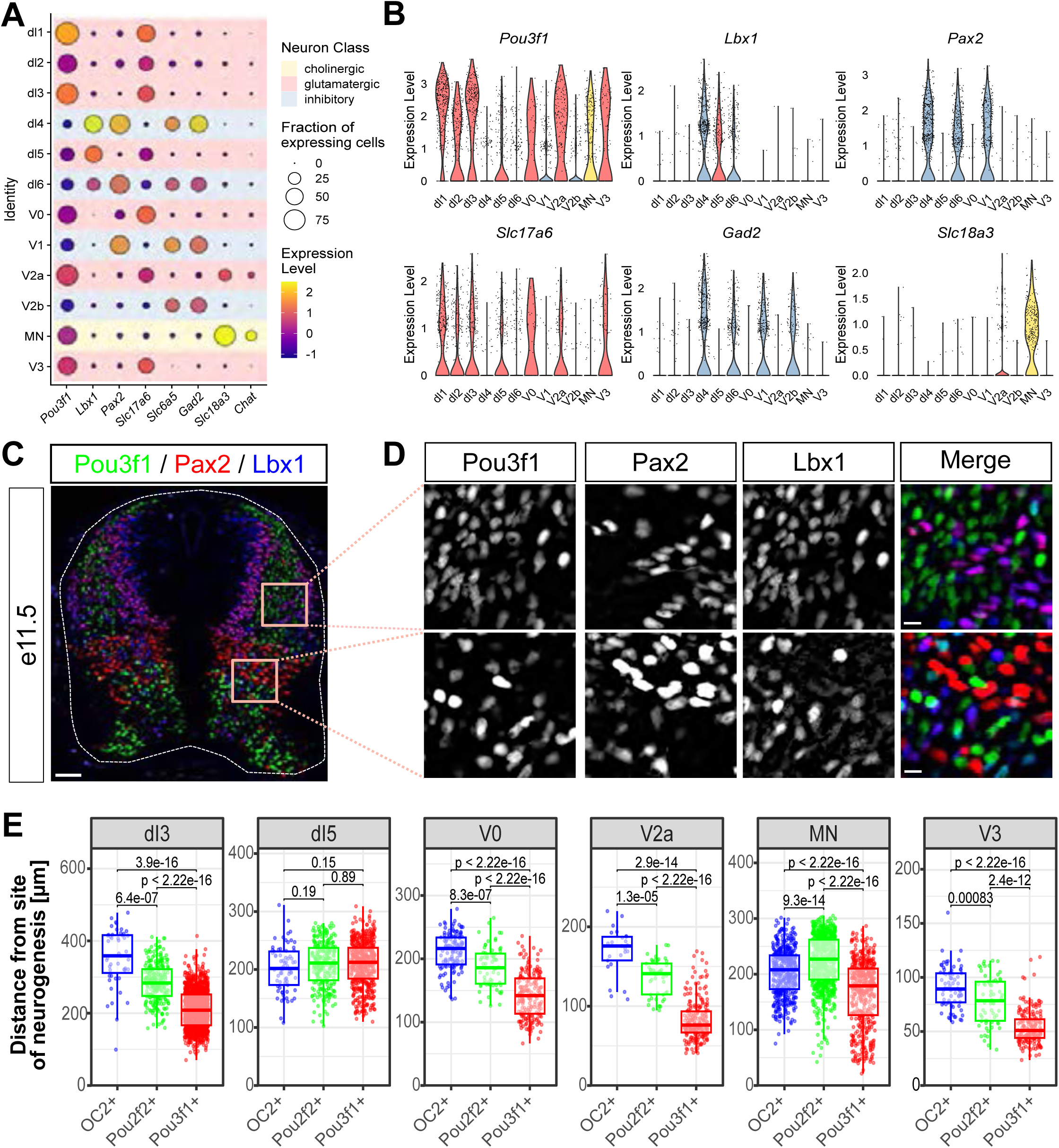
Pou3f1 is not expressed in inhibitory neurons (Related to Figure 2) (A, B) Dot plot (A) and violin plots (B) showing expression of *Pou3f1* and the marker genes *Lbx1* (intermediate dorsal), *Pax2* (inhibitory dI4, dI6 and V1), *Slc17a6* (excitatory), *Slc6a5* and *Gad2* (both inhibitory) and *Slc18a3* and *Chat* (both cholinergic) in e11.5 scRNAseq data from Delile et al. 2019. (C, D) e11.5 spinal cord section stained for Pou3f1, Pax2 and Lbx1. (E) Boxplots indicating the distance of OC2+, Pou2f2+ and Pou3f1+ subsets of different DV classes of spinal cord neurons to their site of neurogenesis. OC2+ neurons are typically further away from the site of neurogenesis than Pou2f2+ neurons, which in turn are further away than Pou3f1+. Scale bars in C = 50 µm; in D = 10 µm

**Figure S5:**
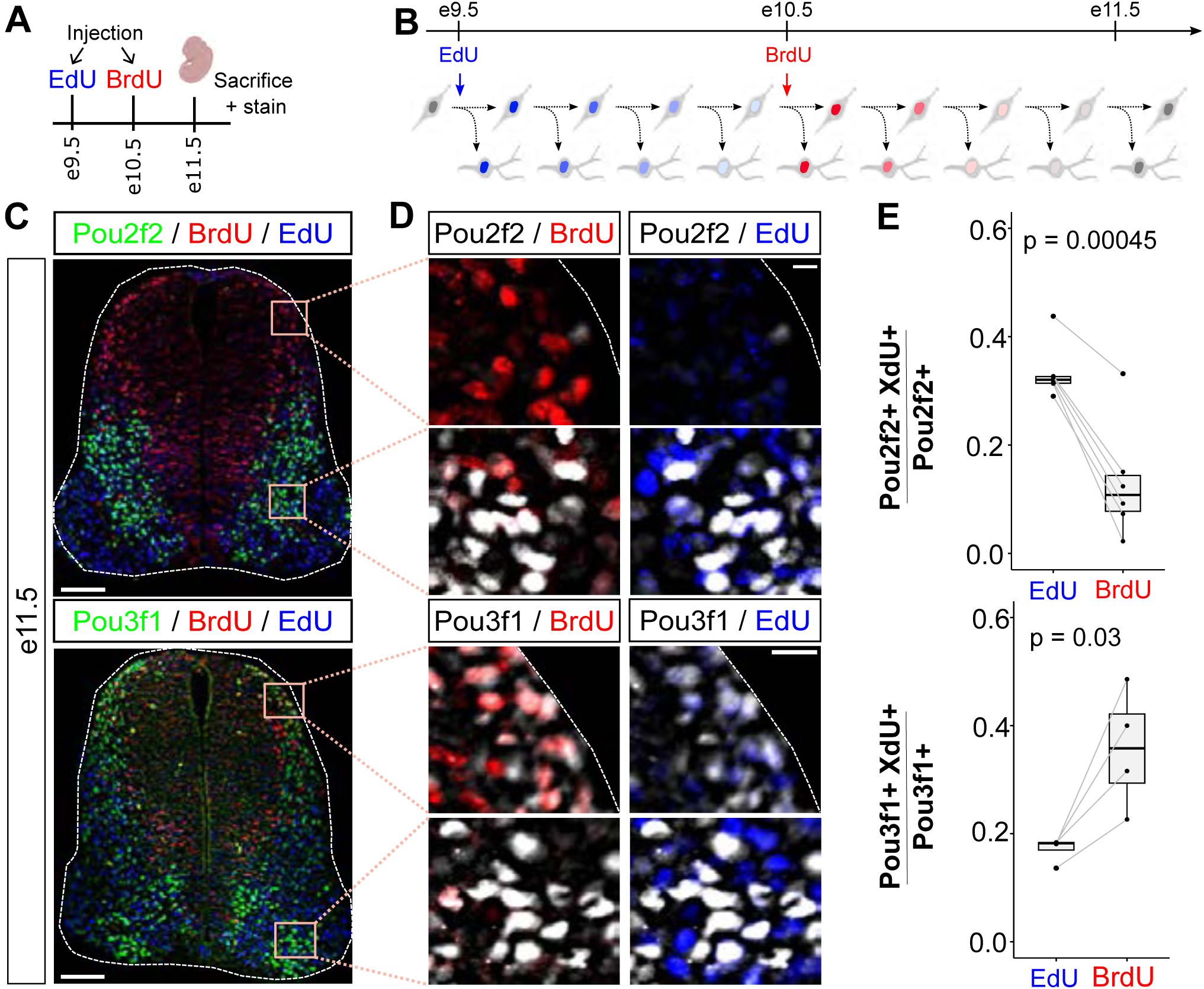
Sequential birth-dating with EdU and BrdU confirms the sequential generation of Pou2f2 and Pou3f1-positive neurons. (A, B) Scheme outlining EdU-BrdU double-birth-dating experiments. Pregnant mothers were injected with EdU at e9.5 and BrdU at e10.5 and incorporation into Pou2f2+ and Pou3f1+ neuronal subsets was analysed at e11.5. (C, D) Spinal cord sections stained for Pou2f2, EdU and BrdU (top) and Pou3f1, EdU and BrdU (bottom). (E) Quantification of the fraction of Pou2f2 (top) and Pou3f1 (bottom) positive neurons labelled by EdU and BrdU (n Pou2f2 = 6 sections from 5 embryos; n Pou3f1 = 4 sections from 3 embryos). Scale bars in C = 100 µm; in D = 10 µm

**Figure S6:**
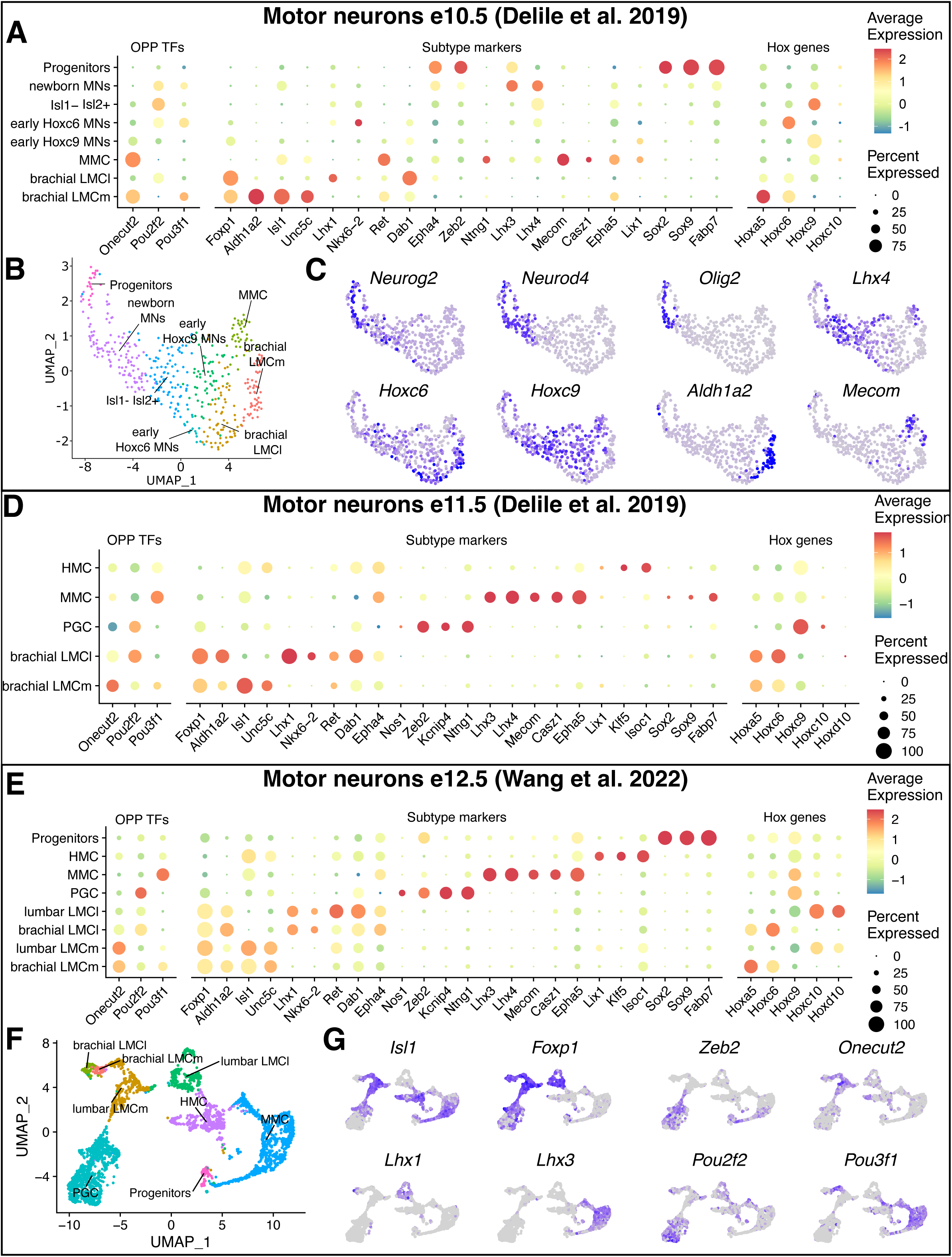
OPP TFs are expressed in different motor columns (Related to Figure 4) (A - C) Analysis of e10.5 MN scRNAseq data from Delile et al. 2019. (A) Dot plot indicating OPP (left), motor column markers (middle) and Hox gene expression (right). (B) UMAP representation color coded for MN subtype identities. (C) Expression of additional marker genes (compare to Fig. 4B). (D) Dot plot indicating OPP (left), motor column markers (middle) and Hox gene expression (right) in MNs at e11.5. (E - G) Analysis of e12.5 MN scRNAseq data from Wang et al. 2022. (E) Dot plot indicating OPP (left), motor column markers (middle) and Hox gene expression (right). (F) UMAP representation color coded for MN column identities. (G) Expression of *OC2*, *Pou2f2* and *Pou3f1* relative to the motor column markers *Isl1*, *Foxp1*, *Lhx1*, *Lhx3* and *Zeb2*.

**Figure S7:**
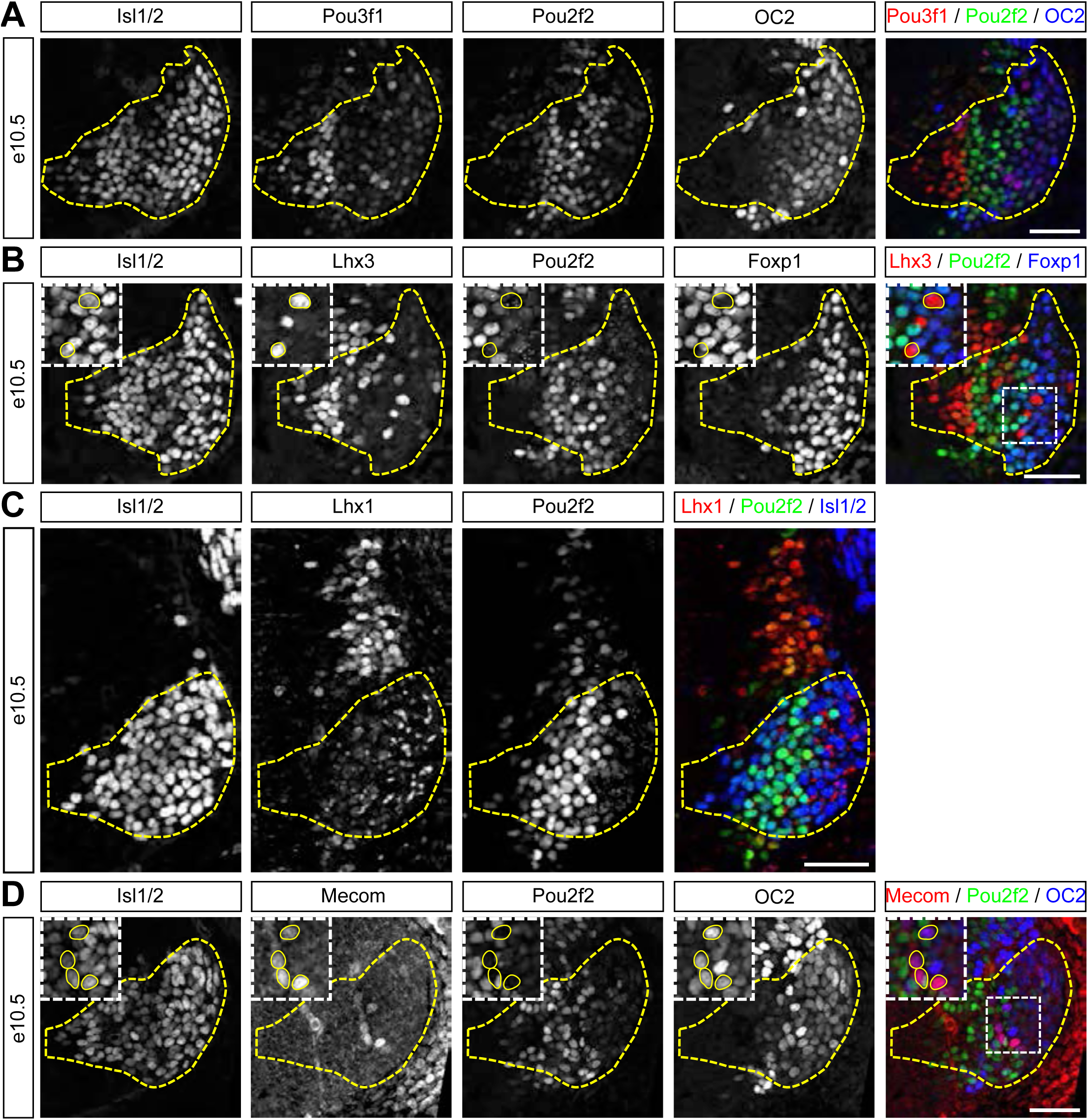
Characterization of OPP expression in e10.5 motor neurons (Related to Figure 4) (A) OPP expression in MNs at e10.5 (single channel views of Fig. 4C). (B) Expression of Pou2f2 relative to Isl1/2, MMC marker Lhx3 and LMC marker Foxp1 (single channel views of Fig. 4D). (C) Downregulation of Isl1/2 in the Pou2f2 stripe precedes high-level Lhx1 expression in MNs (single channel views of Fig. 4F). (D) Early occurrence of a small population of Isl1/2+ Mecom+ OC2+ MMC neurons (single channel views of Fig. 4F). Scale bars = 50 µm

**Figure S8:**
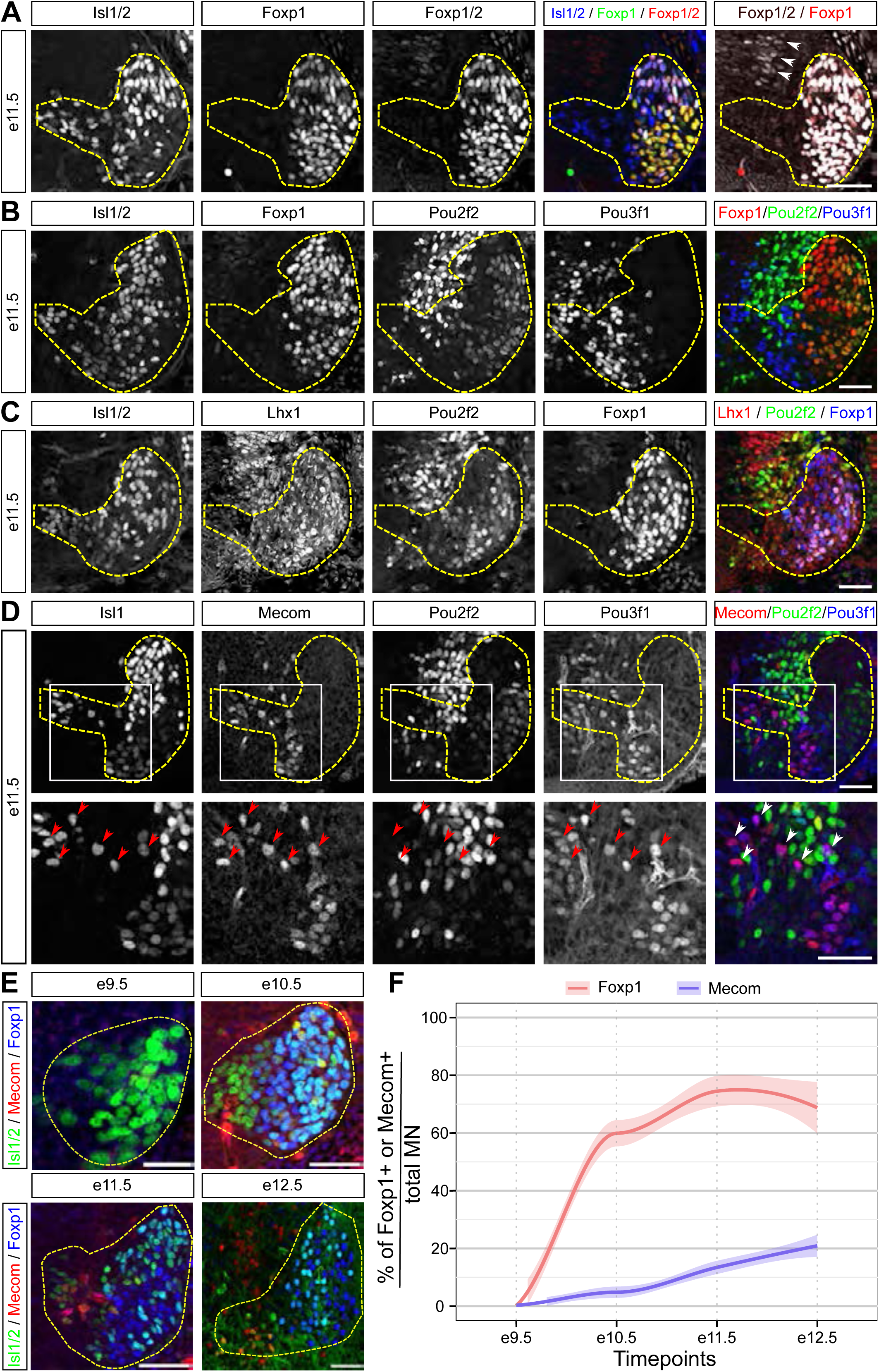
Characterization of OPP expression in e11.5 motor neurons (Related to Figure 4) (A) e11.5 section stained for Isl1/2 and goat anti-Foxp1 (AF4534) and rabbit anti-Foxp2 (ab16046) antibodies. The Foxp2 antibody gave a strong signal in MNs, which normally do not express Foxp2 (Dasen et al., 2008), and a comparably much weaker signal in a more dorsal population, presumably V1 neurons (white arrowheads), suggesting the antibody detects Foxp1 and Foxp2. Contrast for Foxp1/2 and Foxp1 in the last panel was enhanced compared to the other panels to improve the visibility of the signal in the more dorsal cell population. (B, C) Pou2f2 is expressed in LMCl neurons (Isl1/2-negative Foxp1-positive) and Pou3f1 in MMC neurons (Foxp1-negative) (B; single channel views of Fig. 4L). Consistent with these observations, Pou2f2 and Lhx1 stainings coincide in Foxp1+ LMC neurons (C; single channel views of Figure 4M). (D) Pou3f1 is expressed in Isl1, Mecom double-positive MMC neurons (single channel views of Fig. 4N). Many of these Isl1, Mecom, Pou3f1 triple-positive neurons still migrate laterally from the pMN to the MMC (arrowheads). (E, F) e9.5 to e12.5 time-course of LMC and MMC neuron formation. (E) MNs stained for Isl1/2, LMC marker Foxp1 and MMC marker Mecom. (F) Quantification of the percentage of MNs expressing the respective markers. Fraction of Foxp1+ neurons increases between e9.5 and e10.5 and stays afterwards constant, while the fraction of Mecom+ neurons only begins to increase after e10.5 (n e9.5, e10.5 = 23 sections from 6 embryos, n e11.5 = 17 sections from 5 embryos, n e12.5 = 6 sections from 3 embryos). Scale bars in A, B, C, D, E = 50 µm; in E (e9.5) = 25 µm

**Figure S9:**
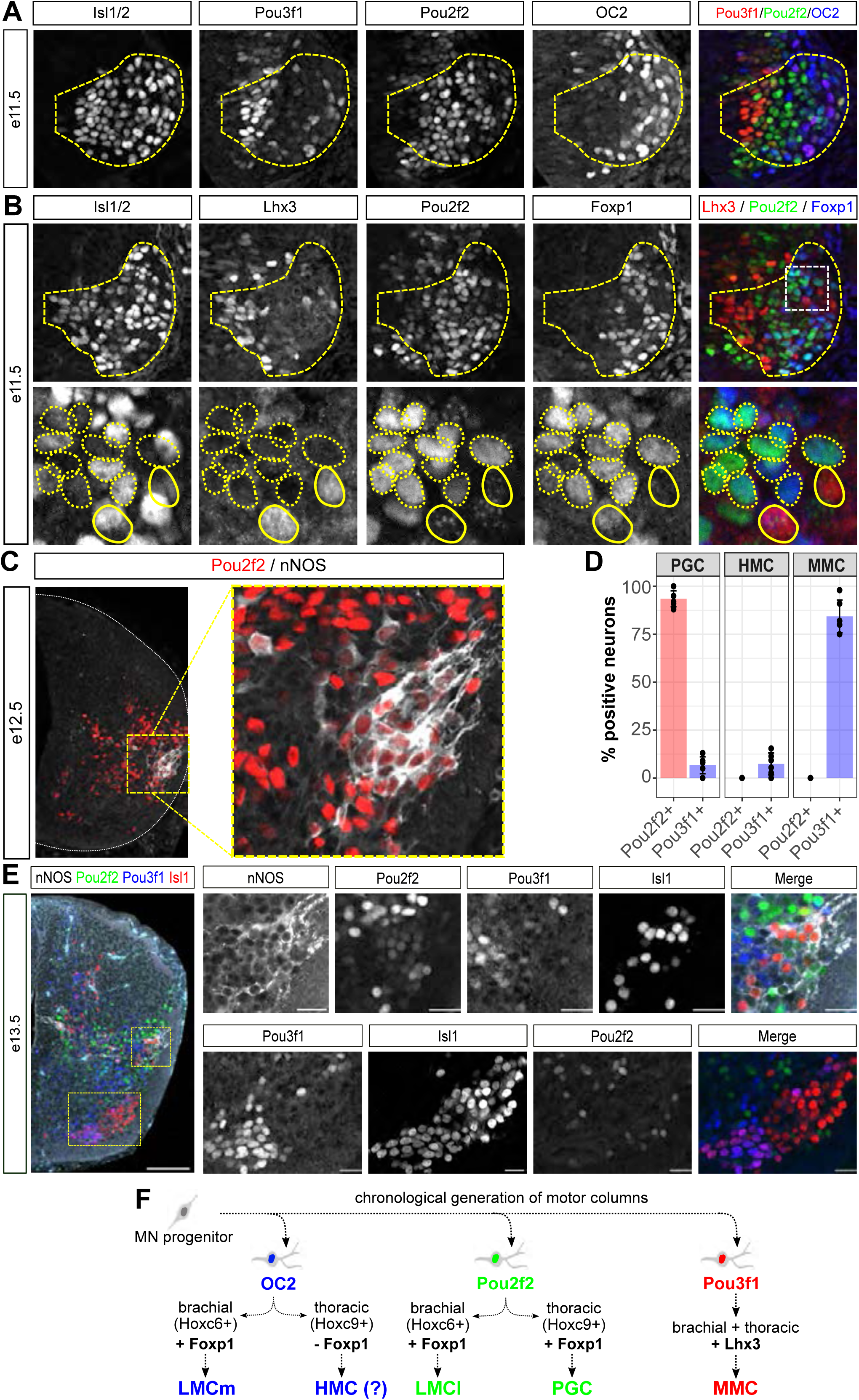
Characterization of OPP expression in thoracic MNs (Related to Figure 4) (A) e11.5 thoracic MNs stained for OC2, Pou2f2, Pou3f1 and the MN marker Isl1/2. (B) e11.5 thoracic MNs stained for Isl1/2, Lhx3, Pou2f2 and Foxp1. Pou2f2 is co-expressed with Foxp1, suggesting Pou2f2+ neurons form the PGC. (C) Co-expression between the Pou2f2 and the PGC marker nNOS in e12.5 thoracic MNs. (D) Quantification of the percentage of Pou2f2+ and Pou3f1+ neurons in the PGC, HMC and MMC at e13.5 (n = 6 sections from 3 embryos). (E) e13.5 thoracic spinal cord section stained for Pou2f2, Pou3f1, the PGC marker nNOS and the MN marker Isl1. (F) Proposed model for the temporally segregated generation of motor columns at different axial levels. Pou2f2+ MNs form the brachial LMCl and thoracic PGC. Pou3f1 is expressed in MMC neurons at both axial levels. OC2 is expressed and functionally required for the formation of the LMCm (Roy et al. 2012). Thoracic OC2+ neurons may contribute to the HMC. Scale bars in E = 50 µm and 25 µm

**Figure S10:**
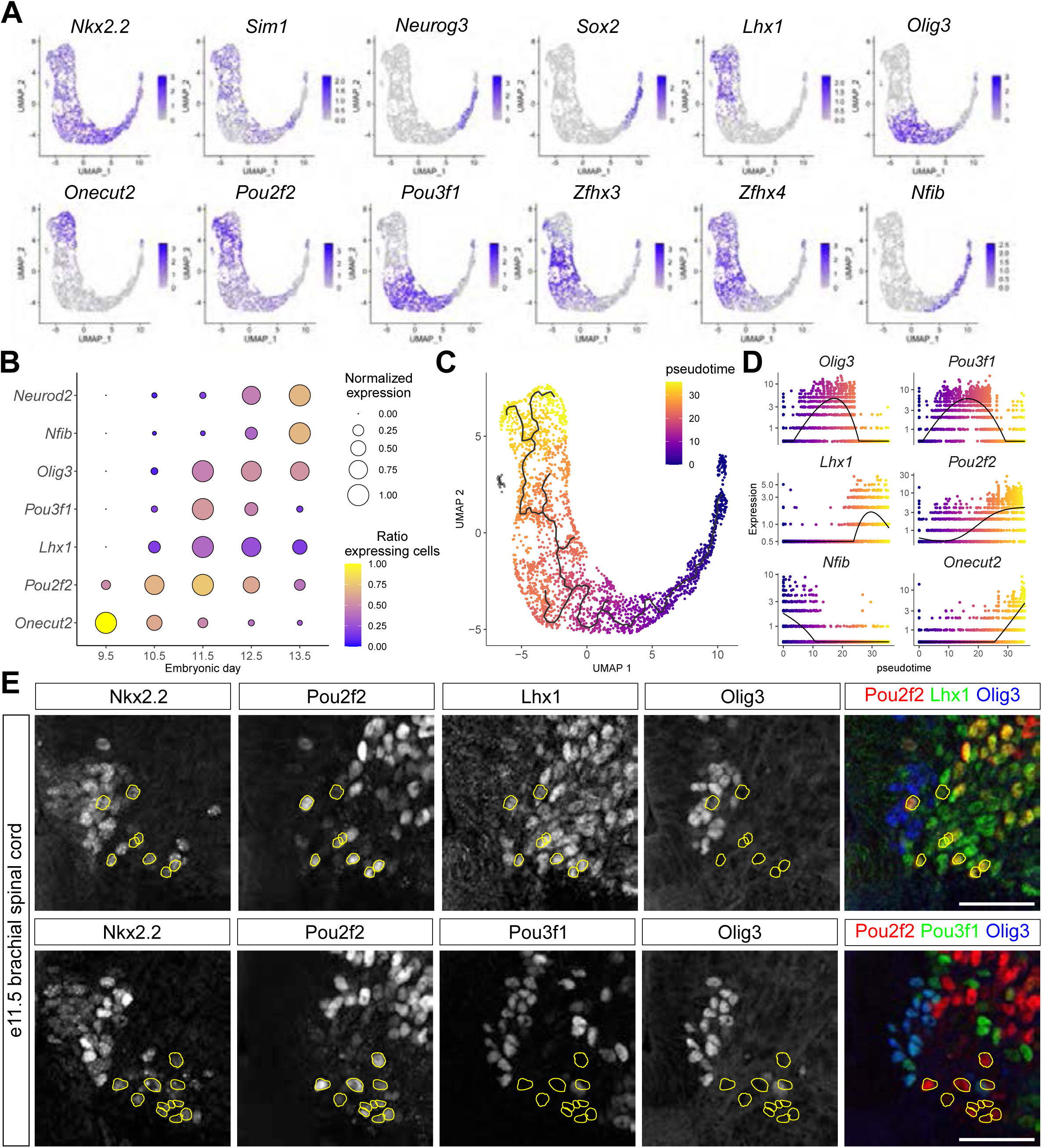
OPP expression in V3 neurons. (A) Expression of V3 (*Nkx2.2, Sim1, Neurog3*), progenitor (*Sox2*), V3_D_ (*Lhx1*), V3_V_ (*Olig3*) and temporal TFs (*OC2*, *Pou2f2*, *Pou3f1*, *Zfhx3*, *Zfhx4*, *Nfib*) markers in V3 neurons in e12.5 scRNAseq data from Wang et al. 2022. *OC2* and *Pou2f2* are expressed in *Lhx1*+ V3_D_ neurons, *Pou3f1* in *Olig3*+ V3_V_ neurons. (B) scRNAseq expression dynamics of temporal TFs, *Lhx1* and *Olig3* in V3 neurons from e9.5 to e13.5. (C, D) Pseudotime gene expression dynamics of the indicated markers based on e12.5 V3 neuron scRNAseq data from Wang et al. 2022. (E) e11.5 brachial spinal cord section stained for V3 neuron marker Nkx2.2, Pou2f2, V3_D_ marker Lhx1 and V3_V_ marker Olig3 (top row) or Nkx2.2, Pou2f2, Pou3f1 and Olig3. Yellow outlines label Nkx2.2 Pou2f2 double-positive cells. Pou2f2 expression coincides with Lhx1 but not Olig3 expression in V3 neurons. Pou3f1 is expressed in Olig3+ V3 neurons. Scale bars in E = 50 µm

**Figure S11:**
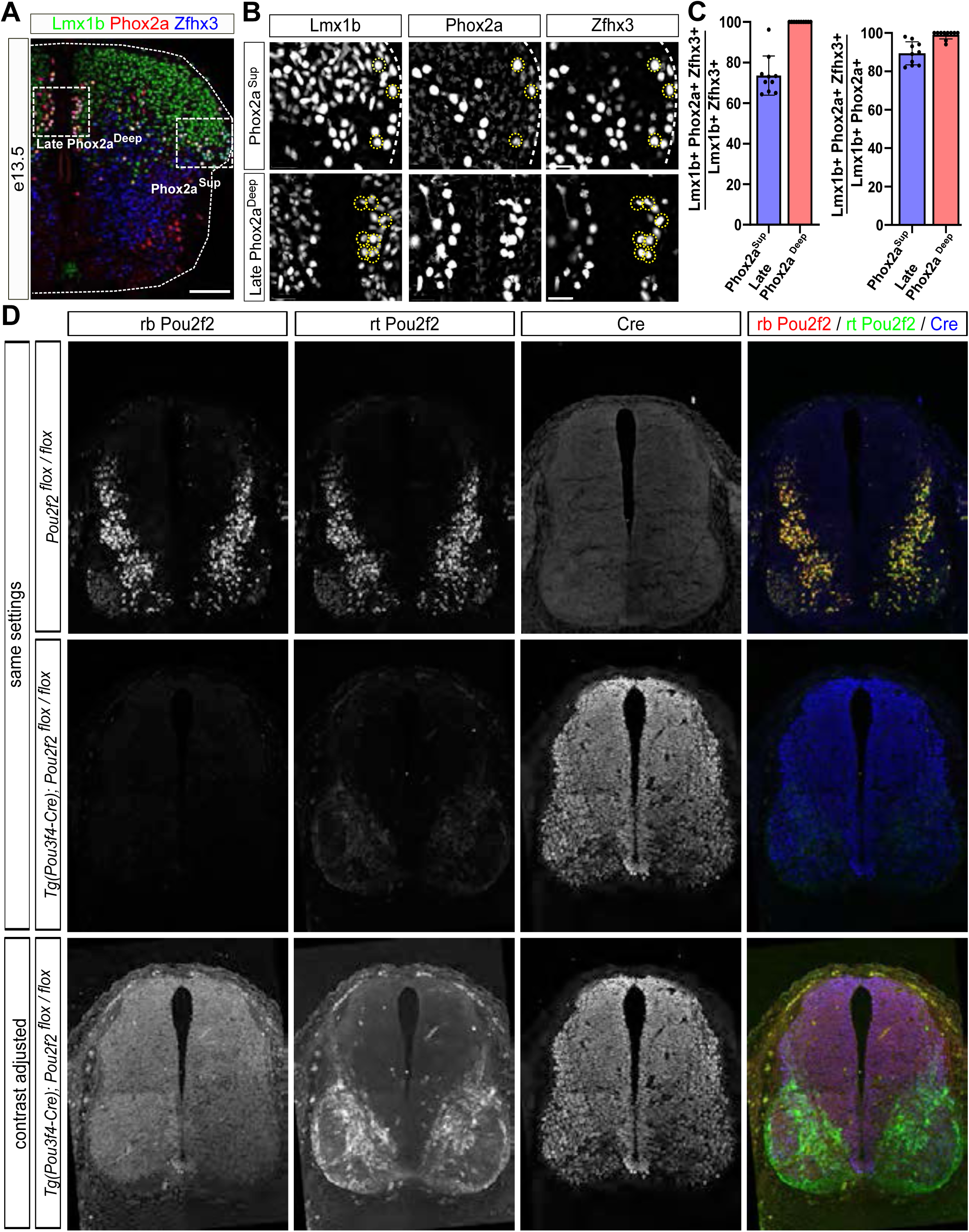
Co-expression of Phox2a and Zfhx3 in Lmx1b+ AS neurons and characterization of Pou2f2 antibodies (Related to Figure 5) (A - C) Phox2a and Zfhx3 expression in the brachial spinal cord at e13.5. (A) Spinal cord section stained for Phox2a, Zfhx3 and the dI5, dILB marker Lmx1b. (B) Higher magnification views of medial and lateral areas of the spinal cord, where Phox2a^SUP^ and Phox2a^DeepLate^ cells reside at this stage. (C) Quantification of Lmx1b+ Zfhx3+ neurons that also express Phox2a (left) and Lmx1b+ Phox2a+ neurons that also express Zfhx3 (right) reveals good agreement between both markers in medial and lateral AS neuron populations (n = 10 sections from 4 embryos). (D) Characterization of Pou2f2 antibodies. e11.5 sections of control (*Pou2f2 flox/flox*; top row) and mutant (*Tg(Pou3f4-Cre) / +; Pou2f2 flox/flox*); middle and bottom row) were stained with rabbit and rat anti-Pou2f2 and anti-Cre antibodies. In control sections, both Pou2f2 antibodies stain the nuclei of the same cells and antibody signal is strongly reduced in the mutants, confirming the specificity of both antibodies. By enhancing the contrast of the mutant sections, residual Pou2f2 cytoplasmic signal can be detected in a similar pattern as endogenous Pou2f2 staining in the wildtype sections with the rat anti-Pou2f2 antibody, consistent with this antibody being raised against amino acids 1-40 of Pou2f2 that should not be affected by the conditional deletion of Pou2f2. With the rabbit anti-Pou2f2 antibody, no specific signal was detected in the mutant sections. Scale bars in A,D = 100 µm; in B = 25 µm

**Figure S12:**
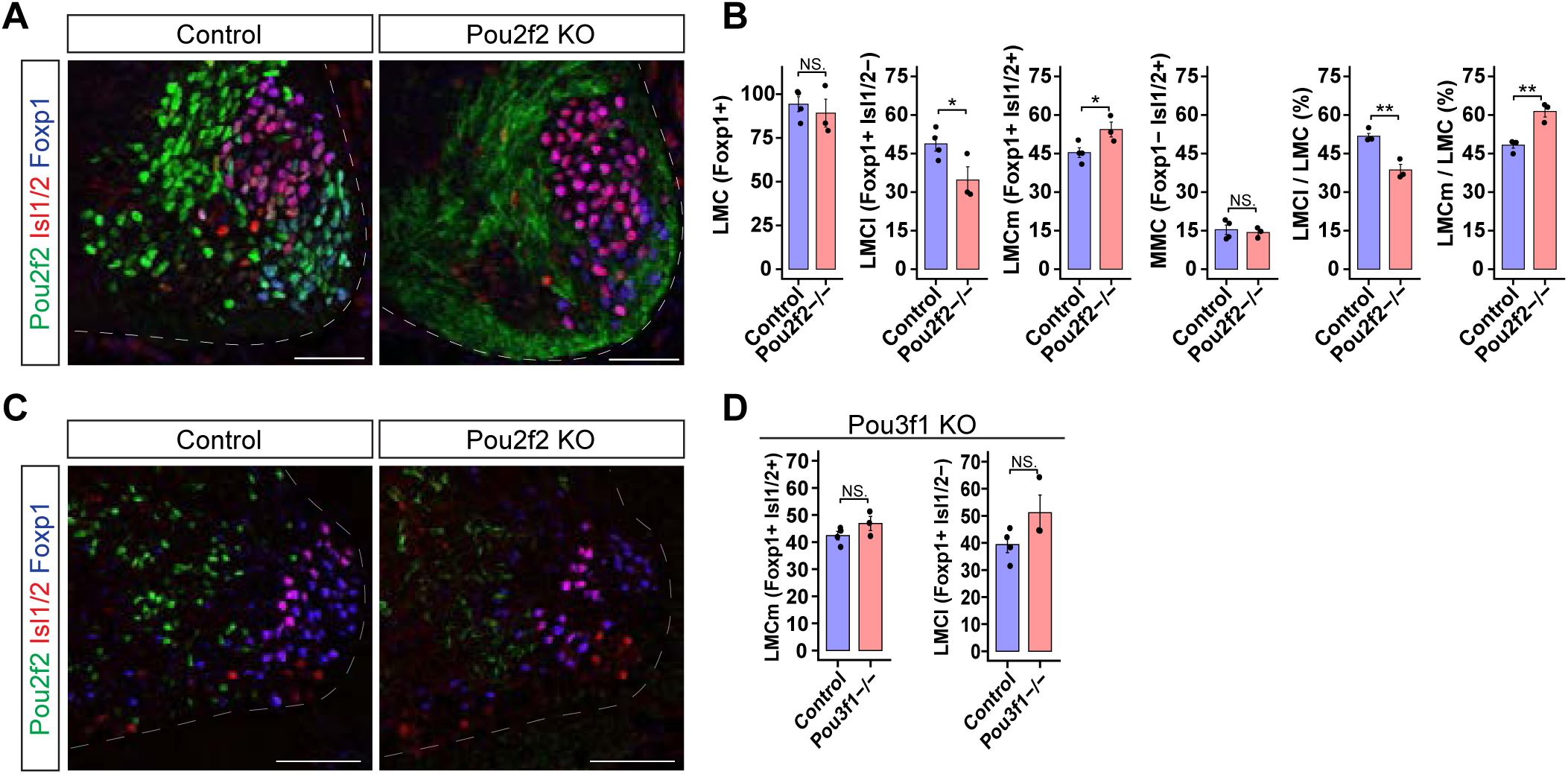
Additional characterisation of motor column specification phenotypes in Pou2f2 and Pou3f1 mutants (Related to Figure 6) (A, B) Staining for Pou2f2, Isl1/2, and Foxp1 in Pou2f2 mutant and control embryos at e11.5. We observed a significant reduction in the LMCl as defined as Foxp1-positive and Isl1/2-negative and a corresponding increase in the LMCm as defined as Foxp1 and Isl1/2 double positive. (C) Staining at e15.5 for Pou2f2, Isl1/2, and Foxp1 in Pou2f2 mutant and control embryos showing the presence of putative LMCl neurons remain by e15.5. (D) Additional quantification related to Fig. 6A indicates no significant increase in either LMCl or LMCm in Pou3f1 mutants. In B and D, Student’s t-test; *p<0.05, **p < 0.01; ***p < 0.001. All data represented as mean ± SEM. Scale bars in A and C = 100 μm.

**Figure S13:**
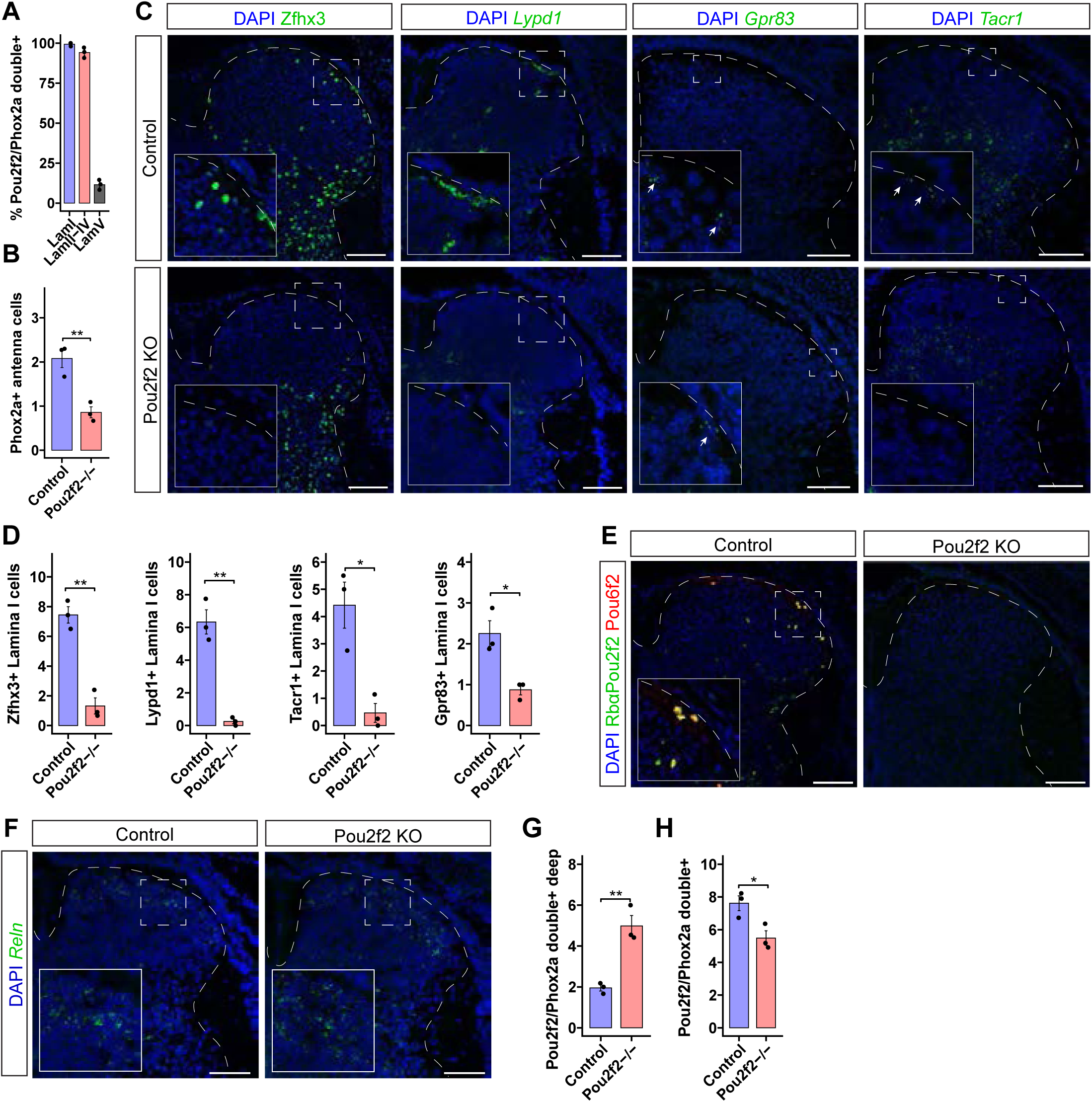
Further characterization of AS subtype specification phenotype in Pou2f2 KOs. (Related to Figure 7) (A,B) Additional quantification related to Fig. 7A. (A) Percent of Phox2a AS neurons that are positive for Pou2f2 by laminae. (B) Number of putative antenna neurons per hemisection (defined as Phox2a AS neurons located between laminae II-IV) is significantly reduced in Pou2f2 KOs compared to controls. (C,D) (C) Expression of various markers of superficial AS neurons in Pou2f2 KOs and controls. (D) Quantification of C showing a significant reduction in all markers of superficial AS neurons assessed. (E) Staining for Pou6f2 and Pou2f2 using the rabbit anti-Pou2f2 antibody used in this study (Abcam Cat#178679, see key resources table). Notably, the Pou6f2 staining is largely restricted to Pou2f2 positive nuclei and is absent in the Pou2f2 KOs. (F) RNAscope ISH for Reln, which is enriched in laminae I and II and is still present in Pou2f2 KOs. (G,H) Additional quantification related to Fig. 7A. (G) Total number of Pou2f2 positive Phox2a neurons per hemisection in Pou2f2 KOs and controls. (H) There is a significant increase in deep Phox2a AS neurons that are labelled by the rat anti-Pou2f2 antibody (RRID:AB_2737828, see key resources table) in the Pou2f2 KOs (i.e. the truncated form of the protein) as compared to controls. In B, D, G, and H, Student’s t-test; *p<0.05, **p < 0.01; ***p < 0.001. All data represented as mean ± SEM. Scale bars in A, C, D = 100 μm.

**Figure S14:**
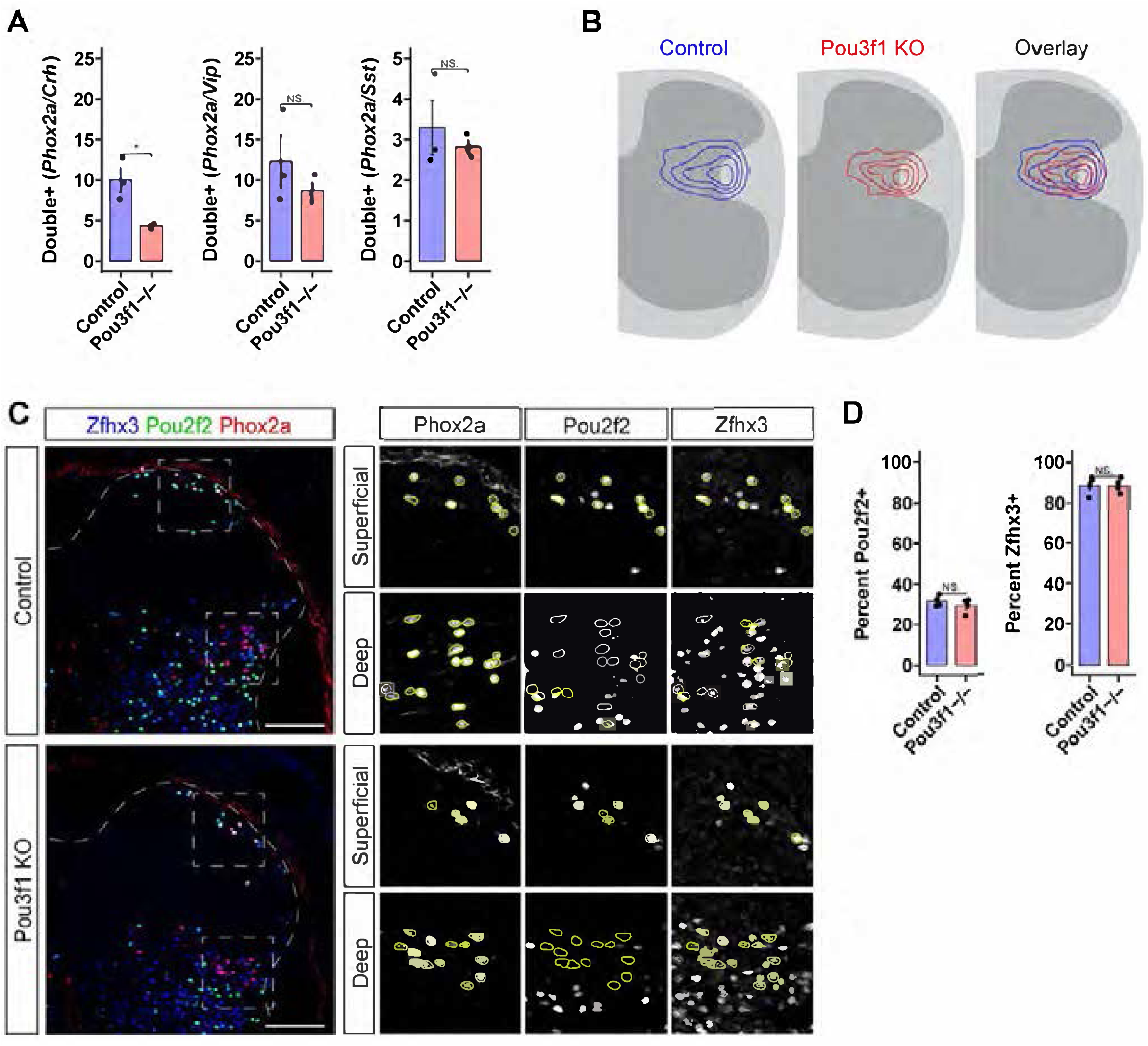
Further characterization of AS subtype specification phenotype in Pou3f1 KOs. (Related to Figure 8) (A, B) Expression of Zfhx3 and Pou2f2 is unchanged in Pou3f1 mutants. (C) Density plots of deep Phox2a neuron location in Pou3f1 mutants and controls showing no change in the distribution of deep Phox2a neurons (n = 4 mutants and controls, Hotelling T2 test p = 0.8973). (D) Absolute number of Phox2a neurons double positive for Crh, Vip, or Sst in Pou3f1 mutants and controls. In B and D, Student’s t-test; *p<0.05, **p < 0.01; ***p < 0.001. All data represented as mean ± SEM. Scale bar in A = 100 μm.

