## Supplementary material for "Evidence for chronological diversification of spinal neuron subtypes by a shared sequence of transcription factors": Key resources table

| **Reagent type (species) or resource** | **Designation** | **Source or reference** | **Identifier** | **Additional information** |
| --- | --- | --- | --- | --- |
| Mice | B6.FVB-Tg(EIIa-cre)C5379Lmgd/J | Jackson Labs | Jax: 003724, RRID:IMSR_JAX:003724 | E2a-Cre |
| Mice | B6;129-Pou2f2^tm1c(EUCOMM)Hmgu/^SdtJ | Louis M. Staudt Laboratory^1^ | Jax: 029132, RRID:IMSR_JAX:029132 | Pou2f2^fl^ |
| Mice | Pou3f1^tm1Mejr^ | Dies Meijer Laboratory^2,3^ | MGI:1857995, RRID:MGI:5515748 | Pou3f1^-^ |
| Antibody | Anti-Foxp1 (Guinea pig polyclonal) | Ben Novitch Laboratory^4^ | Bennett Novitch; UCLA Cat# MI 585, RRID:AB_2811723 | IHC (1:5000) |
| Antibody | Anti-Foxp1 (Goat polyclonal) | R&D Systems | (R&D Systems Cat# AF4534, RRID:AB_2107102) | IHC (1:500) |
| Antibody | Anti-Mecom (Evi1) (Rabbit monoclonal) | Cell Signaling Technology | #2593S, RRID: N/A | IHC (1:1000) |
| Antibody | Anti-Lhx1/5 (Mouse monoclonal) | DSHB | DSHB Cat# 4F2, RRID:AB_531784 | IHC (1:100) |
| Antibody | Anti-Lim1/Lhx1 (Rabbit polyclonal) | Abcam | (Abcam Cat# ab229474, RRID:AB_2924798) | IHC (1:2000) |
| Antibody | Anti-Isl1/2 (Rabbit polyclonal) | Samuel L Pfaff Laboratory^5,6^ | RRID: N/A | IHC (1:20000) |
| Antibody | Anti-Isl1/2 (Mouse monoclonal) | DSHB | (DSHB Cat# 39.4D5, RRID:AB_2314683) | IHC (1:100) |
| Antibody | Anti-Isl1 (Goat polyclonal) | R&D Systems | (R&D Systems Cat# AF1837, RRID:AB_2126324) | IHC (1:1000) |
| Antibody | Anti-nNos (Rabbit Polyclonal) | ImmunoStar | ImmunoStar Cat# 24431, RRID:AB_572255 | IHC (1:1000) |
| Antibody | Anti-nNos (Guinea pig polyclonal) | Synaptic systems | (Synaptic Systems Cat# 432 005, RRID:AB_2832223) | IHC (1:500) |
| Antibody | Anti-Pou2f2 (Oct-2) (Rat monoclonal) | BD Biosciences | BD Biosciences Cat# 562837, RRID:AB_2737828 | IHC (1:1000) |
| Antibody | Anti-Pou2f2 (Oct-2) (Rabbit polyclonal) | Abcam | Cat# 178679, RRID: N/A | IHC (1:1000) |
| Antibody | Anti-Phox2a (Rabbit polyclonal) | Jean-François Brunet (École Normale Supérieure, Paris, France)^7^ | RRID:  AB_2315159 | IHC (1:10000) |
| Antibody | Anti-Zfhx3 (Sheep polyclonal) | R&D Systems | R&D Systems Cat# AF7384, RRID:AB_11127859 | IHC (1:1000) |
| Antibody | Anti-Pou6f2 (Rat polyclonal) | Jay Bikoff (Thomas Jessell Laboratory,  HHMI Columbia University, New York,  United States) | Cat# CU1796 RRID: AB_2665427 | IHC (1:2000) |
| Antibody | Anti-Pou6f2 (Guinea pig polyclonal) | Jay Bikoff (Thomas Jessell Laboratory,  HHMI Columbia University, New York,  United States) |  | IHC (1:10000) |
| Antibody | Anti-Pou3f1 (Oct6) (Mouse monoclonal) | EMD Millipore | MABN738, RRID: N/A | IHC (1:500) |
| Antibody | Anti-Pou3f1 (Oct6) (Rabbit polyclonal) | Abcam | (Abcam Cat# ab272925, RRID:AB_2927579) | IHC (1:500) |
| Antibody | Anti-Pou3f2 (Brn2) (Rabbit monoclonal) | Cell Signaling Technology | #12137, RRID: N/A | IHC (1:1000) |
| Antibody | Anti-Onecut2 (Sheep polyclonal) | R&D Systems | R&D Systems Cat# AF6294, RRID:AB_10640365 | IHC (1:500) |
| Antibody | Anti-Lbx1 (Guinea pig polyclonal) | Thomas Müller (Carmen Birchmeier lab, MDC, Berlin, Germany) | (C. Birchmeier - Max Delbruck Center for Molecular Medicine, Berlin, Germany Cat# Guinea pig anti-mouse Lbx1 polyclonal antibody, RRID:AB_2532144) | IHC (1:2000) |
| Antibody | Anti-Lmx1b (Guinea-pig polyclonal) | Thomas Müller (Carmen Birchmeier lab, MDC, Berlin, Germany) | (T. Müller and C. Birchmeier, Max Delbruck Center for Molecular Medicine; Berlin; Germany Cat# Lmx1b, RRID:AB_2314752) | IHC (1:1000) |
| Antibody | Anti-Vsx2 (Chx10) (Mouse monoclonal) | Santa Cruz Biotechnology | (Santa Cruz Biotechnology Cat# sc-365519, RRID:AB_10842442) | IHC (1:200) |
| Antibody | Anti-Nkx2.2 (Mouse monoclonal) | DSHB | (DSHB Cat# 74.5A5, RRID:AB_531794) | IHC (1:100) |
| Antibody | Anti-Evx1 (Mouse monoclonal) | DSHB | (DSHB Cat# 99.1-3A2, RRID:AB_2246711) | IHC (1:3000) |
| Antibody | Anti-Lhx3 (Rabbit polyclonal) | Biotechne, Novus | (Novus Cat# NBP3-13352, RRID:AB_3588217) | IHC (1:500) |
| Antibody | Anti-Foxp2 (Rabbit monoclonal) | Abcam | (Abcam Cat# ab16046, RRID:AB_2107107) | IHC (1:2500) |
| Antibody | Anti-Neurog2 (Mouse monoclonal) | R&D Systems | (R&D Systems Cat# MAB3314, RRID:AB_2149520) | xx |
| Antibody | Anti-Pax2 (Goat polyclonal) | R&D Systems | (R&D Systems Cat# AF3364, RRID:AB_10889828) | IHC (1:1000) |
| Antibody | Anti-Cre-recombinase (Guinea pig polyclonal) | Synaptic Systems | (Synaptic Systems Cat# 257 004, RRID:AB_2782969) | IHC (1:500) |
| Antibody | Anti-BrdU (rat monoclonal) | Abcam | (Abcam Cat# ab6326, RRID:AB_305426) | IHC (1:500) |
| RNAscope probe | Mm-Lypd1-C1 | ACDbio | Cat #318361, RRID: N/A |  |
| RNAscope probe | Mm-Tacr1-C1 | ACDbio | Cat #410351, RRID: N/A |  |
| RNAscope probe | Mm-Gpr83-C1 | ACDbio | Cat #317431, RRID: N/A |  |
| RNAscope probe | Mm-Crh-C1 | ACDbio | Cat #316091, RRID: N/A |  |
| RNAscope probe | Mm-Vip-C1 | ACDbio | Cat #415961, RRID: N/A |  |
| RNAscope probe | Mm-Sst-C1 | ACDbio | Cat #404631, RRID: N/A |  |
| RNAscope probe | Mm-Pou2f2-O1-C2 | ACDbio | Cat #1170721-C2, RRID: N/A |  |
| RNAscope probe | Mm-Phox2a-C3 | ACDbio | Cat #520371-C3, RRID: N/A |  |
| RNAscope probe | Mm-Reln-C3 | ACDbio | #405981-C3, RRID: N/A |  |
| Chemical | DAPI | Fisher Scientific | #D1306 |  |
| Chemical | Alexa647 Click-iT EdU Imaging  Kit | Thermo Fisher Scientific | Invitrogen #C10340 |  |
| Chemical | BrdU | Sigma Aldrich | Sigma  #B5002 |  |

**Key Resource Table References:**
